# Piglets vocally express the anticipation of (pseudo)-social contexts in their grunts

**DOI:** 10.1101/2020.03.25.007401

**Authors:** A.S. Villain, A. Hazard, M. Danglot, C. Guérin, A. Boissy, C. Tallet

## Abstract

Emotions not only arise in reaction to an eliciting event but also while anticipating it, making this context a way to assess the emotional value of events. Anticipatory studies have poorly considered vocalisations whereas they carry information about the emotional state. We studied the grunts of piglets that anticipated two (pseudo)social events known to elicit positive emotions more or less intense: arrival of a familiar conspecifics and arrival of a familiar human. Both time and spectral features of the vocal expression of piglets differed according to the emotional context. Piglets produced low-frequency grunts at a higher rate when anticipating conspecifics compared to human. Spectral noise increased when piglets expected conspecifics, whereas the duration and frequency range increased when expecting a human. When the arrival of conspecifics was delayed grunts duration increased, while when the arrival of the human was delayed spectral parameters were comparable to those during isolation. This shows that vocal expressions in piglets during anticipation are specific to the expected reward and to the time duration between signal and the delivery of the reward. Vocal expression (time and spectral features) is thus a good way to explore emotional state of piglets during anticipation of challenging events.

## Introduction

Animal behaviour is driven by the motivational system^1^. Animal’s emotions are important feedback mechanisms for modulating the activity of this system. One way to assess the emotional value of an event is to measure the anticipatory activity before the event. Indeed, emotions not only arise in reaction to the challenging event, but also during anticipation of this expected event^2^. Anticipation is goal directed and occurs during the appetitive phase of behaviour^3^, before the consummatory phase. Anticipatory behaviour towards a positive event is adaptive since it is associated with the motivational system that directs the animal from an aversive state (e.g. hungry) to a reinforcing state (e.g. food acquisition; see Spruijt *et al.*^1^). It is suggested that promoting positive anticipation is a way to enhance the quality of life of animals that are under the responsibility of humans^4^.

During anticipation of a positive event, animals are motivated for the event that will arise, and are thus more likely to pay attention to stimuli that are signalling the event itself^3^. Anticipation has been used to evaluate the sensitivity and motivation to different events supposed to be positive like food reward, social contact, and play^3,5–7^. It may also be used to evaluate cognitive judgement bias that are consecutive to a long terms emotional experience^8^.

Anticipation of social contact and play is expressed by an increase of the time spent in a compartment where the given social reward is expected to arrive in rats^5^, and an increase of activity in rats, silver foxes, pigs, horses and lambs^5,7,9–11^. Comparing the behavioural anticipation responses to different events allows evaluating both the relative valence and the intensity of the emotion associated to the expected events. For instance, lambs express a higher elevation of the activity and more behavioural transitions before food than before play, suggesting that food reward is a more intense positive event^7^. In hens, behavioural anticipation is different according to the quality of the food reward^6^. In other cases, the increase of the level of activity is not specific to the quality of the anticipated event. For instance, in a study in pigs, the level of locomotor activity during anticipation of a positive (food) and a negative (frightening event) events do not differ^12^. In silver foxes, differences in anticipatory behaviour before different food rewards are shown in the posture of the ears, but not in the level of activity^10^. This suggests anticipatory behaviour may be event-specific and species-specific e.g.^13^. Thus specific anticipatory behavioural activations, i.e. activity and spatial location, may not be sufficient to highlight differences in anticipatory behaviour in all cases.

In addition, vocalisations may be interesting measures to explore emotional content of the anticipation phase. Indeed, vocalisations have an emotional content in many species ^14^. However, vocalisations have been poorly included in the anticipatory behaviour ethograms until now. One exception are the rats’ frequency modulated ultrasonic vocalisations (50kHz)^15^. In horses, low-pitched vocalisations (i.e. nickers) are proposed by Peters *et al.*^11^ as expressions of positive anticipation but the authors were not able to score them in their study. In pigs, high-frequency vocalisation are suggested to be a good indicator of the emotional state during anticipation of different events^12^. The probability that a pig makes high-frequency vocalisations is higher before negative than positive events^12^. Pig are a good candidate to study vocalisations during anticipation because the variability in vocal expressions according to emotions has been reported both according to the valence and the arousal dimensions of emotions^16–19^. Thus vocalisations quality, especially of grunts, would be a good indicator to evaluate anticipation in pigs.

In the present study, we wanted to measure the vocal expression in piglets during the anticipation of events with different emotional values. In the farming context, pigs may experience various kinds of (pseudo)social events. Pigs being social animals the presence of familiar conspecifics has a highly positive valence^20^. Pigs also experience interactions with humans and may develop positive relationship with them after a period of positive reinforcing interactions, such as brushing and calmly speaking, compared to control animals^21–23^. Positive anticipation of human contact is possible in captive non domestic animals^24^, and we tested this in pigs.

The aim of this study was to test whether piglets reared in-group could vocally express anticipation of arrival of social partners and arrival of a familiar human caregiver and if vocalisations were different according to the social characteristics of the reward. We first conditioned piglets to associate a visual and acoustic signal to the arrival of familiar conspecifics and another signal to the arrival of a familiar human caregiver. Half of the piglets had previously received additional positive contacts with the human prior to the conditioning, leading to two groups of piglets with different degrees of familiarity toward the human. To complete the investigation of emotional values of the anticipated event, we carried out a final test delaying the arrival of the expected partner. We measured both their behavioural and vocal activity, as well as the acoustic structure of their grunts, during the signal, e.g. the anticipatory phase, and then after the signal when the reunion was delayed. We hypothesized that expecting familiar conspecifics has a positive valence and induces a high arousal state for all piglets, compared to expecting a familiar human, even considering the quality of his familiarity. If vocal signals reflect emotional states, we expect them to have a different signature when anticipating conspecifics compared to a human. If having received additional contacts with the human prior to the conditioning modifies the emotional state of piglets, we expect a different anticipatory vocal signature between groups that had or had not received additional care.

## Results

### Anticipatory behaviour differs between piglets expecting the arrival of a Human or conspecifics

One trial was separated in five phases, tested as factors: before the signal was broadcasted (phase −1), while the signal was broadcasted, i.e during the anticipation phase (phase 0) and when the arrival of the partner was delayed of 1.5 minutes, which was segmented in three 30-second phases (1, 2 and 3).

Behavioural parameters were used to build two scores using a multivariate analysis carried out in two steps. A linear discriminant analysis was first computed on a subset of data containing the first two phases of the trial, e.g. before (phase −1) and during the signal (=phase 0, or anticipation phase). Two behavioural scores corresponding to the first two linear discriminant functions (LD1 and LD2) were thus built. On the remaining dataset, containing the last three phases (after the signal: named 1, 2, and 3), a projection was computed on LD1 and LD2, allowing to test for differences between the phases in the 2D behavioural space.

The first behavioural score (LD1) was negatively correlated with the time spent near the upcoming partner’s door and the time spent watching this door and positively correlated with the number of zones explored (table 1). Statistics showed a significant interaction between the phase of the trial and the partner (X^2^_4_ = 13.9, p = 0.008, figure 1A, significance letter from a to d). During trials of anticipation of the human partner, the signal led to a significant decrease of LD1 compared to the initial phase (H partner, phase −1 *vs.* 0, T.ratio = 5.97, p < 0.001). After the signal, while the arrival of the partner was delayed, LD1 increased and then remained stable (H partner, phase 0 *vs*. 1: T.ratio = −1.64, p = 0.83, 0 *vs.* 2: T.ratio = −3.88, p = 0.004, 0 *vs.* 3: T.ratio = −3.82, p = 0.005); remaining at the same level as before the signal (H partner, phase −1 *vs.* 1:2:3, |T.ratio| < 2.62, p > 0.21). During trials of anticipation of conspecifics, the signal led to a significant decrease of LD1 compared to the initial phase (C partner, phase −1 *vs.* 0, T.ratio = 7.33, p < 0.001). After the signal, LD1 did not change while the arrival of the partner was delayed (C partner, phase 0 *vs.* 1:2:3, |T.ratio| < 2.7, p > 0.18), but tended/were to be different than during the initial phase (C partner, phase −1 vs. 1, T.ratio = 2.5, p = 0.25; phase −1 vs. 2, T.ratio = 3.1, p = 0.06; phase −1 vs. 3, T.ratio = 5.1, p <0.001). Prior to any signal, LD1 differed depending on the type of partner (phase −1, C *vs.* H, T.ratio = −14.74, p < 0.001). No interaction between the phase of the trial and the additional care treatment (H versus H+) was found (X^2^_4_ = 0.69, p = 0.95) and no main effect of the treatment was found (X^2^_1_=0.13, p = 0.72).

**Table 1:** Loadings of linear discriminant functions 1 and 2 (respectively LD1 and LD2) of the behavioural analysis of anticipation in piglets trained to expect the arrival of familiar conspecifics or a familiar human. In lines, the parameters used to build the functions and in columns their respective loadings on the first two functions. Note that this table concerns the data from phase ‘−1’ (before the signal) and phase ‘0’ (during the signal) and that a factor taking into account the partner, the phase and the treatment was used to discriminates the groups. These loading were then used to project the data gathered during the violation of expectations test (trial 12) and statistics were run on LD1 and LD2 after projection as two behavioural scores.

| Behavioural scores | LD1 | LD2 |
| --- | --- | --- |
| Total time spent freezing | 0.062 | -0.181 |
| Averaged duration spent per zone | -0.044 | 0.438 |
| Number of zones explored | 0.226 | 0.134 |
| Total time spent watching upcoming partner's door | -0.224 | 0.356 |
| Number of time watching upcoming partner's door | -0.188 | 0.664 |
| Total time spent in zones near upcoming partner's door | -1.083 | -0.708 |

**Figure 1:**
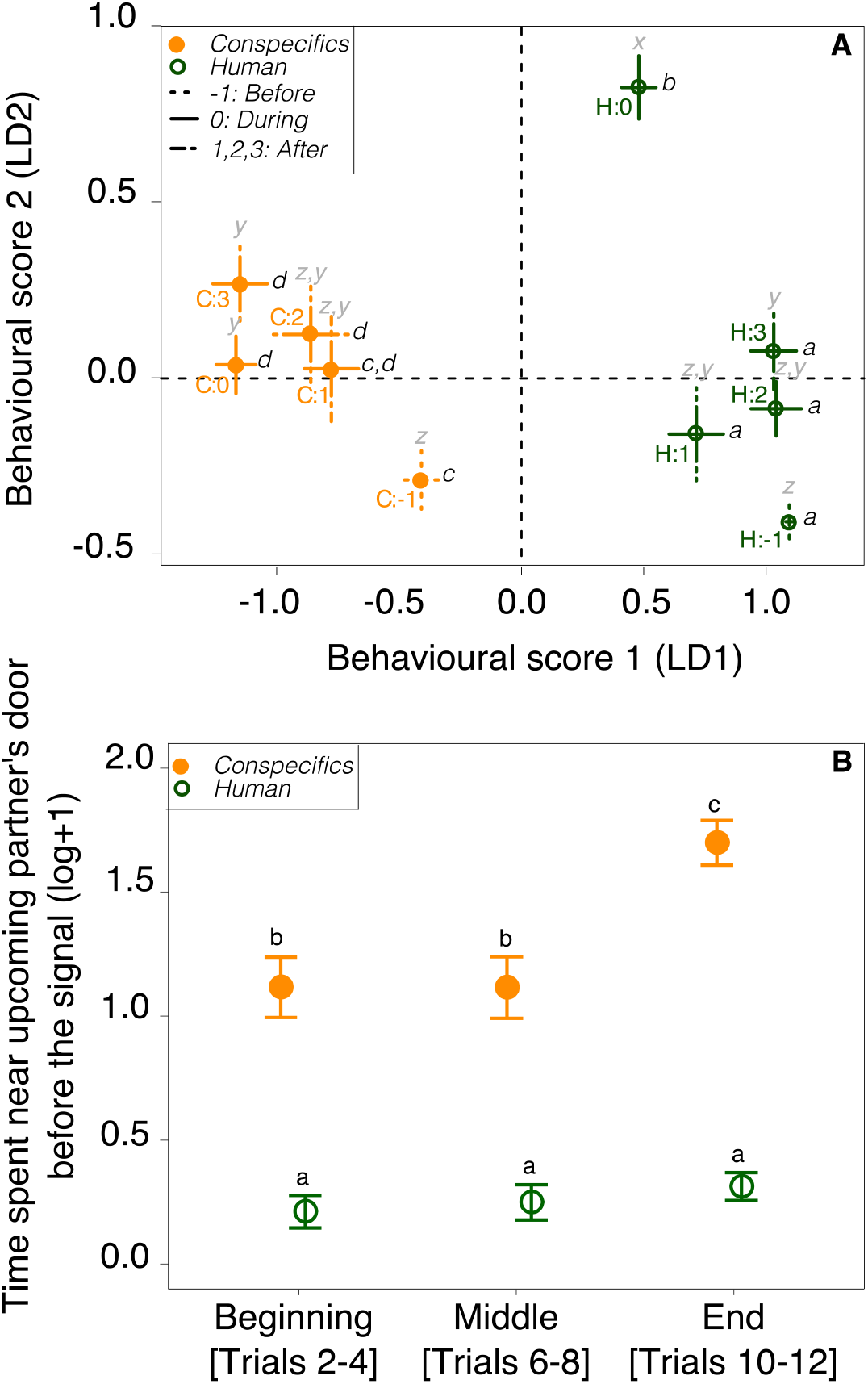
Behavioural responses of piglets to the anticipation of the entrance familiar conspecifics (C, filled orange points) or a familiar human (H, empty green points). Points and bars represent mean ± se per group. A: behavioural space with LD1 and LD2 behavioural scores showing the significant interaction between the phase (−1, 0, 1, 2 and 3) of the test and the type of partner. Phase −1 corresponds to the time before the broadcasting of a signal, phase 0 corresponds to the time during the signal and phases 1, 2, and 3 are 30 second segments during the violation of expectation period (90 seconds in total). Letter from a to d and x to z show significant differences on LD1 and LD2 respectively. B: Time spent near upcoming partner’s door before the broadcasting of any signal (phase-1 only) along the conditioning (grouping trials to create three factors: beginning, middle and end). Letters shows significant differences between groups. All model anova tests, estimates and pairwise post hoc tests with Tukey contrasts are available in tables S1 S2 and S3 respectively as supplementary material.

The second behavioural score (LD2) was negatively correlated with the time spent near the upcoming partner’s door and the time spent freezing and positively correlated with the time spent watching the upcoming partner’s door, the number of times watching the upcoming partner’s door and time spent per zone (table 1). Statistics showed a significant interaction between the phase of the trial and the partner (X^2^_4_ = 49.1, p < 0.001, figure A1, significance letter from x to z). During trials of anticipation of the human partner, the signal lead to a significant increase of LD2 compared to the initial phase (H partner, phase −1 *vs*. 0, T.ratio = 11.29, p < 0.001). After the signal, LD2 significantly decreased while the arrival of the partner was delayed (phase 1) and then remained stable (H partner, phase 0 *vs.* 1:2:3 |T.ratio| < 6.37, p < 0.001), at the same level as before the signal (H partner, phase −1 *vs.* 1:2:3, |T.ratio| < 3.20, p > 0.05). Such effects were not significant for trials of anticipation of conspecifics, for which only a trend was found toward an increase of LD2 during the anticipation phase (C partner, phase −1 *vs.* 0, T.ratio = −3.01, p = 0.08). No difference was found between the anticipation phase and the phase while the arrival of the partner was delayed (C partner, phase 0 vs. 1:2:3 |T.ratio| < 1.48, p > 0.90). Before the signal we found no effect of the upcoming partner on LD2 (phase −1, C *vs.* H, T.ratio = 1.10, p = 0.98). No interaction was found between the phase of the test and the treatment (X^2^_4_ = 2.59, p = 0.63) and no main effect of the treatment was found (X^2^_1_=2.26, p = 0.61; figure 1A, significance letter from x to z).

LD1 and LD2 were mainly explained by the location of the piglet in the experimental room, and piglets significantly spent more time near the conspecifics’ door (see stats for LD1). So differences between LD1 prior to the emission of any signal could be explained by either location biases in the test room or the expression of a preference toward the conspecific door.

We thus tested the effect of the trial number (grouping trials at the beginning, the middle and the end of the experiment, figure 1B) on the time spent near the upcoming partner’s door. If piglets expressed a preference toward the conspecific door along the conditioning, the interaction between the trial number and the partner should be significant. Statistics showed a significant interaction between the partner and the conditioning trial number (X^2^_2_ = 11.96, p = 0.003): although piglets spent more time near the conspecifics’ door than near the human’s door, independently from the on-going partner, piglets increased their time near the conspecifics’ door at the end of the conditioning (C partner, middle vs. end of the conditioning, T.ratio = 4.19, p = 0.001), but did not increase their time spent near the human door (H partner, pairwise tests between all trial factors, |T.ratio| < 0.8, p > 0.97).

For full statistical report, refer to tables S1and S3 of the supplementary material.

### Vocal dynamic differs between piglets excepting a human or conspecifics

The grunt rate, during the anticipation phase (phase 0), was tested using the mean individual inter-grunt interval. The inter-grunt interval was significantly lower during anticipation of conspecifics than during anticipation of human (X^2^_1_=4.35, p = 0.037, figure 2), without any interaction with the treatment (X^2^_1_=0.037, p = 0.85, table S1).

**Figure 2:**
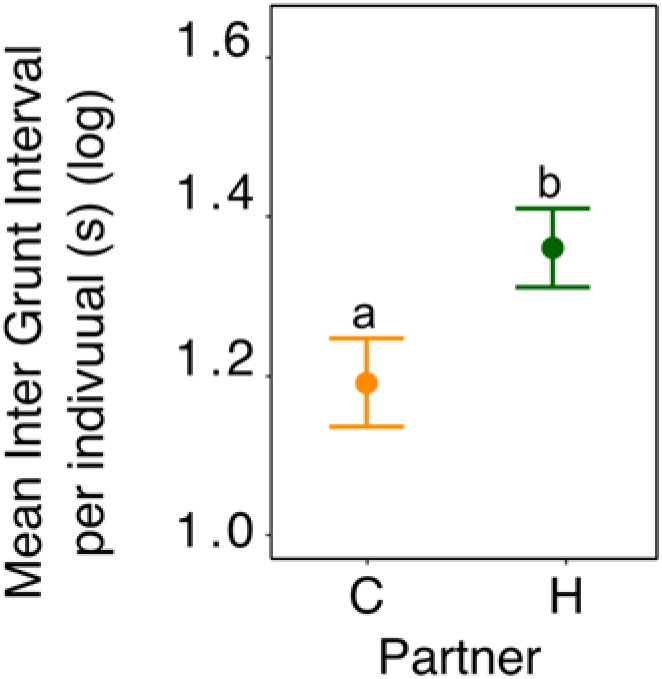
Vocal dynamic in piglets during the anticipation phase (phase 0). Mean ± se inter-grunt interval per individual per type of intra or interspecific partner. Different letters represent significant differences between expected partners. All model anova tests, estimates and pairwise post hoc tests with Tukey contrasts are available in supplementary tables S1, S2 and S3.

### The acoustic structure of anticipatory grunts differs between piglets excepting the arrival of a human or conspecifics, and depends on the degree of familiarity with the human

The acoustic structure of 2270 grunts (see table S4 for data composition) was analysed using the duration of the call and a built-in spectral score, i.e. the first linear discriminant function built from nine acoustic parameters representative of the call spectrum (LD1, table 2). Similarly, to the behavioural analysis, the spectral score LD1 was built using the first two phases of the test (before and during the signal). Then for the last three phase, LD1 values were computed after projection on the same linear discriminant analysis. In order to test for acoustic responses to anticipatory signals, interactions between the phases of the test, the partner and the treatment were tested using linear mixed effect models.

**Table 2:** Loading of the first linear discriminant function (LD1) following the spectral analysis of grunts in piglets. In lines, the parameters used to build the functions and in column their respective loadings on the first function. The linear discriminant analyse was made with the data before and during the signal (phases −1a and 0). A factor taking into account the partner, the phase and the treatment was used to possibly discriminate or not the groups.

| parameters | Loading on the first function |
| --- | --- |
| Mean | -1.442 |
| Median | -0.161 |
| Mode | 0.198 |
| Q25 | 0.034 |
| Q75 | 0.308 |
| Centroid | -1.442 |
| SH | 2.888 |
| SFM | 2.249 |
| Entropy (H) | -2.817 |

#### The acoustic structure of anticipatory grunts differs between piglets excepting a familiar human or conspecifics

To test the effect of the social quality of the partner (conspecifics vs. familiar human), the interaction between the phase of the trial and the type of partner was studied and was significant regarding both the duration of the call and the spectral structure (X^2^_4_ = 50.3, p < 0.001, figure 3A and X^2^_4_ = 63.5, p < 0.001 figure 3D respectively).

**Figure 3:**
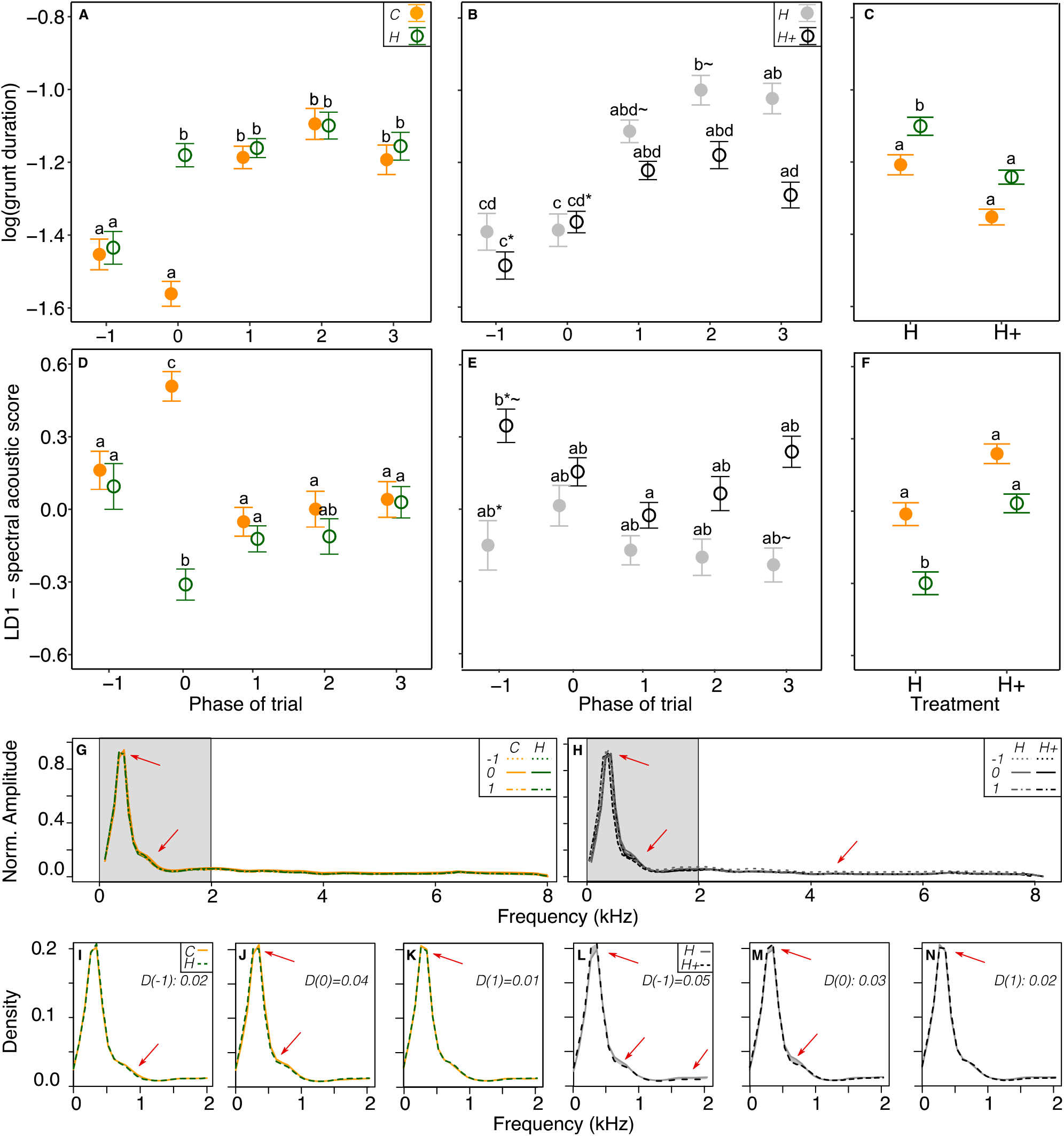
Acoustic structure (mean ± se) of grunts in piglets depending on the type of partner [familiar conspecifics (C, filled orange points) or human (H, empty green points)] and treatment [additional contacts group (H+, empty black points) or minimal contact group (H, filled grey points). A. and B: evolution of grunt duration along phases of the trials. C: grunt duration according to partner and treatment. D and E: evolution of spectral score LD1 along phases of the trials. F: Spectral score LD1 according to partner and treatment. Phases correspond to: before the signal (phase −1), during the signal i.e anticipation phase (phase 0) and after the signal, i.e during the violation of expectation phases (phases 1,2,3 of 30 seconds). Letters shows significant differences between groups. * and ~ symbols identify statistical trends between two groups. All model anova tests, estimates and pairwise post hoc tests with Tukey contrasts are available in tables S1, S2 and S3 respectively as supplementary material. G-N: representation of mean spectra per group (computed with ‘meanspec’ function, ‘seewave R package, wl=512, overlap = 50%). Each line represents a mean over all the grunts extracted in a specific phase and/or treatment/type of partner group. Due to extremely low variability in the spectrum per group, standard errors of the mean of all spectra are not visible on the plots. The number of grunts used per group is available table S4 (56<N<241, median=101 grunts on a total of 2270). Arrows indicate where the changes are the strongest. I-N: zooms in on the 0-2kHz frequency range, for which the coefficient D(phase) correspond to a metric of spectral dissimilarity (0<D<1, computed with ‘diffspec’ function, ‘seewave’ R package).

**Figure 4:**
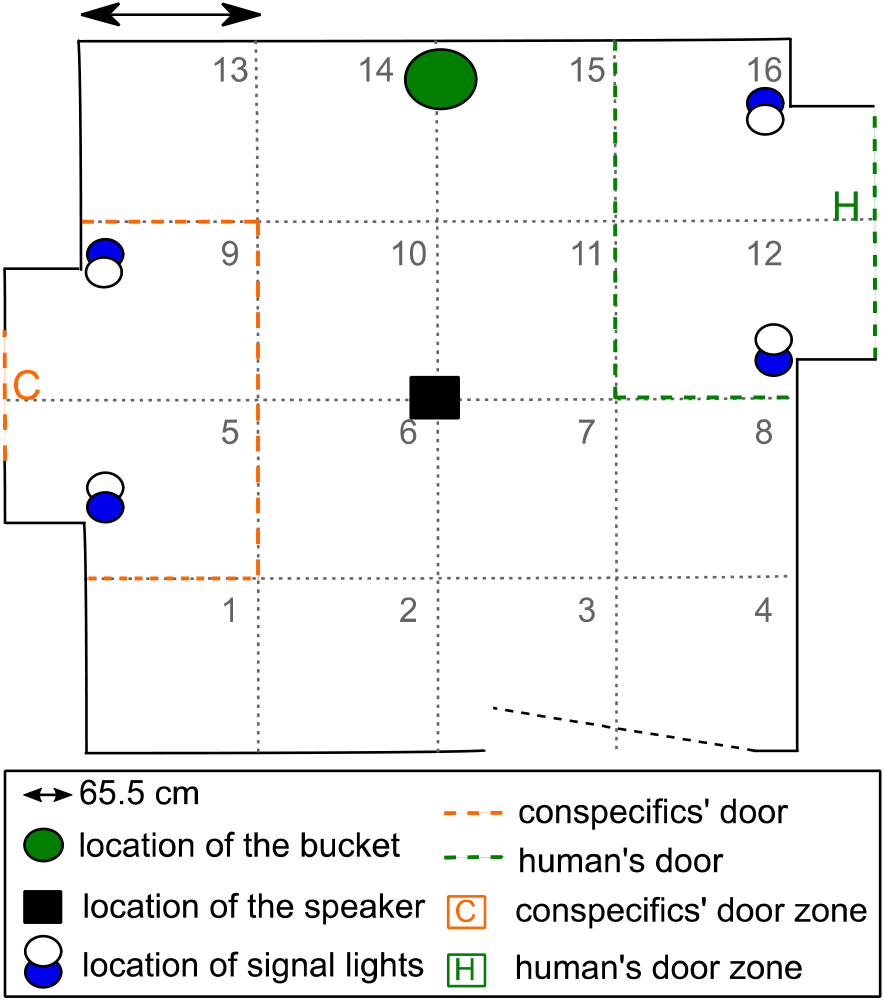
Schematic of the test room. The acoustically isolated room contained three doors: the human’s door (H on the right), the conspecifics’ door (C on the left), and the entrance door (at the bottom), which remained located this way during the entire experiment. A speaker, in the center of the room and at 1m high from the ground broadcasted a 2sec signal, associated to the upcoming partner. Blue and white lights, around the partner’s door were used as visual signal, either blue or white (from two to 20 seconds) announcing the entrance of the partner. When the human entered the room, they would bring a bucket and sit for two minutes bringing additional care to the tested piglet. For behavioural analyses, the room was separated in 16 zones to allow quantifications of mobility and location in the room as well as partner door zones (either C or H).

Considering the duration of the grunts, during trials of anticipation of the human partner, grunt duration was longer than before the signal (partner H, phase −1 *vs.* 0, T.ratio = −4.79, p < 0.001). After the signal, during the phase while the arrival the partner was delayed, grunt duration remained as high as during the anticipatory phase (partner H, phase 0 *vs.* 1:2:3, |T.ratio| < 1.66, p > 0.82) and was higher than before the signal (partner H, phase −1 *vs.* 1:2:3, |T.ratio| > 3.59, p < 0.012). During trials of anticipation of conspecifics, the signal did not affect grunt duration (partner C, phase −1 *vs.* 0, T.ratio = 2.42, p = 0.315). After the signal, grunt duration was higher during all the phases while the partner was delayed (partner C, phase 0 *vs.* 1:2:3, |T.ratio| > 7.32, p < 0.0001). The duration of grunt differed between the type of partner during the anticipation phase (phase 0, C *vs.* H, T.ratio = −8.29, p < 0.001) but were not different before (phase −1, C *vs.* H, T.ratio = −0.14, p = 1.00) nor during the violation of expectation phase (phase 1:2:3, C *vs.* H, |T.ratio| < 0.34, p = 1.00).

Considering the acoustic spectral score LD1 (table 2), statistics showed a significant interaction between the phase of the trial and the type of partner (X^2^_4_ = 63.5, p < 0.001, figure 3D). During trials of anticipation of the human partner LD1 decreased during the anticipation phase compared to the preceding phase (H partner, phase −1 *vs.* 0, T.ratio = 3.95, p = 0.003) and increased after the signal, while the arrival of the partner was delayed (H partner, phase 0 *vs.* 1: T.ratio = −3.74, p = 0.007, phase 0 *vs.* 2: T.ratio = −2.85, p = 0.12, phase 0 *vs* 3: T.ratio = −4.72, p < 0.001) and returned to LD1 values measured before the signal (H partner, phase −1 *vs.* 1:2:3, |T.ratio| < 1.29, p > 0.96). During trials of anticipation of the conspecifics, LD1 increased during the anticipation phase compared to the previous phase (H partner, phase −1 *vs.* 0, T.ratio = −3.97, p = 0.003) and decreased after the signal, while the arrival of the partner was delayed (C partner, phase 0 *vs.* 1:2:3, |T.ratio| > 4.90, p < 0.001) and returned LD1 values measured before the signal (C partner, phase −1 *vs.* 1:2:3,|T.ratio| < 0.66, p = 1.00). LD1 was significantly higher during anticipation of the conspecifics than the human (phase 0, C *vs.* H, T.ratio = 9.00, p < 0.001), and there was no difference between the partner for the other phases (phases −1:1:2:3, C *vs.* H, |T.ratio| < 0.66, p = 1.00).

#### The acoustic structure of anticipatory grunts differs with the degree of familiarity toward the human

To test the effect of the familiarity of the human partner, the interaction between the phase of the test and the treatment was studied and was significant regarding both the duration of the call and the spectral structure (X^2^_4_ = 12.4, p = 0.015 figure 3B, and X^2^_4_ = 14.8, p = 0.005 figure 3E). Regarding grunt duration, in the H+ group (handled piglets), we found no difference in the duration of grunts during the anticipation phase compared to the other (H+ group phase 0 *vs.* −1:1:2:3, |T.ratio| < 2.95, p > 0.094) but grunts were longer after the signal, while the arrival of the partner was delayed, than before the signal (H+ group phase −1 *vs.* 1:2:3, |T.ratio| > 3.75, p < 0.007). In the H group, the duration of grunts increased after the signal, while the arrival of the partner was delayed (group H, phase 0 vs. 1:2:3, |T.ratio| > 3.38, p < 0.025) but did not differ between the anticipation phase and the preceding phase (group H, phase −1 vs. 0, T.ratio = 0.23, p = 0.55). A significant interaction was also found between the type of partner and the treatment, independently of the phase (X^2^_1_=5.84, p = 0.016, figure 3C): piglets from the H group produced longer grunts than piglets from the H+ group during trials of anticipation of a human (H partner, H *vs.* H+, T.ratio = 2.69, p = 0.04) but there was no difference during trials of anticipation of conspecifics (C partner, H *vs.* H+, T.ratio = 1.31, p = 0.56).

Regarding the spectral score LD1, in the H+ group, a significant decrease of LD1 was found from the phase before the signal to the first phase while the arrival of the partner was delayed (H+ group, phase −1 *vs.* 1, T.ratio = 3.26 p = 0.038) and all other comparisons between phases did not differ (H+ group, all other phases, |T.ratio| < 2.89, p > 0.11). In the H group, no difference was found in LD1 between all phases (H group, pairwise between all phases |T.ratio| < 1.80, p > 0.73), Within phases, no difference was found between treatments but LD1 in the H+ group tended to be higher than LD1 in the H group before the signal was broadcasted (phase −1, H vs. H+, T.ratio = −3.06, p = 0.078). All other comparisons between treatments were non-significant (all phases, H vs. H+, |T.ratio| < 2.69, p > 0.19). A significant interaction was found between the type of partner and the treatment, independently from the phase (X^2^_1_=6.45, p = 0.010, figure 3F). LD1 of grunts produced by piglets from the H group in trials of anticipation of the human partner were lower than in trials of anticipation of conspecifics (H group, C vs. H, T.ratio = 4.02, p < 0.001), but not for piglets from the H+ group (H+ group, C vs. H, T.ratio = 1.37, p = 0.52).

To illustrate spectral changes in grunts, mean spectra per group were represented in figure 3, identifying the two types of significant interactions found with the phase of the test: type of partner, (figure 3G, 3I-K) and treatment (3H, 3L-N). Mean spectra representing all frequency range of grunt (0-8 kHz, figure 3G and 3H) were represented as well as zooms in on the specific 0-2 kHz range to identify the changes with arrows (figure 3I to 3N), in the latter zooms in, a coefficient of difference between the mean spectra was calculated to have a better ideas of difference between considered groups.

## Methods

### Ethical note

Experiments were performed under the authorization no. APAFIS#17071-2018101016045373_V3 from the French Ministry of Higher Education, Research and Innovation; and were in agreement with the French and European legislation regarding experiments on animals.

### Subjects and housing conditions

Sixty weaned female piglets (in two replicates), *Sus scrofa domesticus*, bred from crosses between Large White and Landrace females and Piétrain males were used for this study from 28 to 62 days after birth. Animal housing and experiments took place at the experimental unit UEPR (UE 1421, INRAE France).

One piglet was removed in the middle of the experiment due to health issues independent from the experiment. Piglets from the same litter and having similar weight (<1 kg difference) were housed by three in a 1.2 x 1.3m pen on plastic duckboard and panels visually isolated pens. One chain per pen was used for enrichment. Food and water were available *ad libitum*. Artificial lights were turned on from 8:00 to 17:00 and temperature was maintained between 26 and 27 °C. Two identical rooms were used (5 pens per room per replicate).

### Experimental treatment: human additional contacts

Two experimental treatments were generated as follows:

- <u>A group with minimal human contact, H group:</u> Control piglets from 10 rearing pens received the minimal amount of daily contact with a stockperson (a 1.70m high male) required for feeding, cleaning and health checking. The stockperson wore dark green shirt and pans with brown shoes.
- <u>A group with additional human contacts, H+ group:</u> Animals from the 10 other rearing pens received, in addition to daily care given by the stockperson as for H group, 29 sessions of additional human gentle tactile contact from one of the two experimenters (both women, both between 1.70-1.73, balanced number of pens attributed). The experimenters wore the same overalls and boots each time they interacted with the pigs; i.e. blue overalls and dark green boots. The handling procedure, using gentle tactile contacts, was similar to Tallet *et al.*^21^. Those additional contacts were given from the day of weaning until day 39, with three sessions per day (with a two hour break in between) except at weekends. The order of the pen was balanced across days. We confirmed that the additional human contact treatment (H+) induced a positive attraction toward the human in a standard human-piglet reunion test (supplementary material, fig. S1).

### Two-way associative learning and induction of anticipation

Pen piglets were habituated to the test room for 10 minutes, two days before the start. The conditioning took place between day 42 and 62 after weaning, lasted twelve days, with two trials per day with at least three hours between trials of the same day.

#### (Pseudo) social events

All piglets were individually trained to learn to associate two different signals with the arrival of two different (pseudo)-social partners for 2 minutes: either three pen mates (partner = Conspecifics) or a familiar human (partner = Human). When entering the room, the human sat on a bucket and interacted with the piglet, in the same way she interacted with them during the taming phase. For piglets from the H+ group, the human was already familiar (same as the taming phase) whereas for piglets from the H group, the human was unfamiliar and became familiar along the conditioning.

#### Associative learning signals

Associative learning stimuli were chosen to facilitate learning since the aim was not to test learning abilities but the way piglets would anticipate the reunions. One signal announcing the entrance of a partner combined one visual and one auditory stimulus ^25^: visual stimuli were lights (blue or white) lighting on nearby door and auditory stimuli were tones (296 Hz or 3100 Hz, broadcasted from a speaker (Mipro MA-100su, Mipro Electronics Co, Taiwan). Four visio-auditory combinations were thus built and their occurrences were balanced across all experimental piglets.

#### Associative learning trials

Twenty-four trials were run by piglet; 12 with each partner. For each trial, the target piglet entered the experimental room and remained alone between 10 to 30 seconds to avoid habituation to the start of the signal (phase −1) before the signal started and lasted between two and 20 seconds (phase 0, anticipation). After the end of the signal, the partner entered the room. Piglets from the same pen were tested one after the other and the order was balanced from one trial to the other to avoid confounding effect of the order within one pen. The order of the pens was balanced from one day to the other to avoid confounding effect of the period of the day. Piglets were reunited once with each of the possible partner each day (balancing between the morning and the afternoon), except on days 6 and 8 for which they were reunited the same partner in the morning and the afternoon trials to avoid habituation to alternating reunions. This design was inspired by Reimert *et al*.^25^.

#### Inducing and testing anticipation

To generate an anticipatory phase (phase 0) prior to the arrival of the partner, the duration of the signal was gradually increased along the conditioning^25^ [trials 1-3: two seconds, trials 4-5: five seconds, trials 6-7: 10 seconds, trials 8-9: 15 seconds, trials 10-12: 20 seconds]. To allow the recording of the vocalisations produced during the anticipatory phase, only the visual stimulus was prolonged and not the auditory stimulus which was kept at two seconds for all trials. Only trials containing a signal (and thus an anticipatory phase) of 20 seconds were analysed. In order to test for anticipation, we needed to disrupt the associative learning. We thus delayed the entrance of the supposedly expected partner on the last and twelfth trial: the signal stopped but the partner entered only after one and a half minute (divided into three 30s-phases, named phases 1, 2 and 3).

### Behavioural measures

Behaviours were monitored using a camera (Bosh, Box 960H-CDD) and annotated using *The Observer*© *XT 14.0* (Noldus, The Netherlands) software. The squared room was split in 16 equally dimensioned zones to assess the mobility and exploratory behaviour of the piglet: every time the shoulders of the piglet crossed a zone, a zone change was scored. The following behaviours were monitored and standardised per minute for each phase: time spent near conspecifics’ and human’ door zones, time spent watching conspecifics’ and human’ doors, number of time the piglet watched the conspecifics’ and human’ doors, number of zones explored, average time spent per zone, time spent static in the room. Behavioural scores were then calculated to quantify global responses (see below).

### Acoustic measures and analyses

#### Acoustic monitoring

Vocalisations were recorded with a AKG C314 microphone placed in the center of the room and at one-meter-high, connected to a Marantz MD661MK2 recorder. Vocalisations produced during each phase of the trial were manually annotated per vocal type (grunt, squeak, bark, scream and mixed calls), after visual inspection of spectrograms on Praat® software. Only grunts were deeply analysed as they were the most expressed. However, additional observational data on other call types are available in the supplementary document (fig. S1).

#### Acoustic measures on grunts

A spectro-temporal analysis was performed with custom-written codes using the Seewave R package^26^ implemented in R^27^. After a 0.2-8 kHz bandpass filtering (‘fir’ function), a standardised grunt was detected when amplitude crossed a 5% amplitude threshold (‘timer’ function) to measure the duration. After amplitude normalisation, the following spectral parameters were calculated (‘specprop’ function, FFT with Hamming window, window length = 512, overlap = 50%): mean, median, first (Q25) and third (Q75) quartiles, interquartile range (IQR), centroid and mode (all in Hz). The grunt dominant frequency (in kHz) was also calculated (‘dfreq’, 50% overlapping FFTs, window length = 512), which is the mean over the grunt duration of the frequencies of the highest level of energy. Parameters measuring noisiness and entropy of the grunt were: Shannon entropy (sh), Spectral Flatness (Wiener entropy, sfm) and Entropy (H) [combining both Shannon and Temporal envelop entropy, length = 512, Hilbert envelop). Two linear acoustic parameters were used: the logarithm of grunt duration and a built-in spectral acoustic score with all spectral parameters (see below). Table of acoustic data available in supplementary material (table S4).

### Statistical analyses

#### Behavioural and acoustic scores

To assess changes in global behavioural/acoustic responses during anticipation, parameters were used to build scores using multivariate analyses carried out in two steps. First, a linear discriminant analysis was computed on a subset of data containing the first two phases of the test, maximizing differences between groups of an *ad hoc* factor ‘phase:treatment:partner’. Two behavioural scores (LD1 and LD2) and one spectral acoustic score (LD1) were built. On the remaining dataset (trial 12: phases 1, 2, 3 for which the entrance of the partner was delayed), a projection was computed on LDs scores, allowing to test for differences in behavioural/acoustic space(s).

#### Statistical tests and validation

We tested for differences in LDs scores since the question using a delayed entrance of the partner was to know whether the piglets would keep the state, they would have during anticipation, return to the state they had in initial phase or exhibit intermediate response. All statistics were carried out on R^27^. A linear mixed effect model (‘lmer’ function, ‘lme4’ R package) was built to test two-way interactions between factors ‘phase of the trial’ (phases: −1, 0, 1, 2, 3), ‘partner’ (Human or Conspecifics) and ‘treatment’ (additional H+ or minimal human contacts H). The factor ‘replicate’ (first or second) was also tested in interaction with ‘treatment’ and ‘partner’. Piglet’s identity was put as random factor (repeated measures per piglet). This model was used to test for behavioural scores (LD1 and LD2), the spectral acoustic score (LD1) and the duration score (log). For vocal rhythm (inter grunt interval), the model was simplified to the study of the anticipation phase only (phase 0), since the metric calculated highly depended on the number of observations. The following two-way interactions were tested: ‘partner’ and ‘treatment’, ‘replicate’ and ‘partner’, ‘replicate’ and ‘treatment’. To test for biases in the piglet’s location in the room prior to the emission of any signal (phase −1), the time spent near to the upcoming partner’s door (parameter loading the most on the LDs), was used as response variable and trials were grouped in a three-level factor: ‘beginning: trials 2-4’, ‘middle: trials 6-8’ and ‘end: trials 10-12’. The model tested the three-way interaction between ‘trial’, ‘partner’ and ‘treatment’ and two-way interactions between ‘replicate’ and ‘partner’ or ‘treatment’. All linear models were validated by visual inspection of the symmetrical and normal distribution of the residuals (‘plotresid’ in ‘RVAideMemoire’ R package). Anovas were computed on models to test for significant effects of explanatory variables (‘car’ R package). Model estimates and pairwise post hoc tests were computed using Tukey correction for multiple testing (‘lsmeans’ R package). A complete report of statistics is available as supplementary material (tables S1-S3).

## Discussion

The paradigm developed in this study was built to analyse in piglets the acoustic expression of anticipation of (pseudo)-social events: arrival of two pen mates or a familiar human. We tested two degrees of familiarity with a human, created by handling sessions given to half of the piglets proven to induce a positive link with the human.

The behavioural analysis showed that piglets were able to anticipate the social reunion: piglets did show a short-term specific response during the anticipation phase compared to the other phases (approach of zone where partners entered during the signal, attention behaviours toward this location, and specific vocal expression, at least for one of the possible partners). When the arrival of the partner was delayed, the duration of grunts increased for both partner. Longer grunts had already been associated to negative emotional valence^18,19,28^, which confirms that the delay lead in piglets to a negative emotional state, a situation of non-correspondence with expectation, whatever the partner. These results allow us to conclude that we did succeed in generating a specific anticipatory state during the tests. Those changes were not solely due to the signal emission, as some were persistent even after the signal (e.g. spatial position). We thus confirm the cognitive ability of weaned piglets for associative learning, and for developing expectations from their environment^12^.

Piglets expressed different behaviours toward partners, showing a preference for their conspecifics compared to the familiar human, which reflect different emotional values (valence, intensity) of the partners. Piglets spent more time near the area of the room where the conspecifics were supposed to enter along the condition sessions, compared to the area where the human partner was supposed to enter. In addition, during the delay phase, piglets expecting conspecifics expressed reactions similar to the anticipation phase, whereas piglets expecting a human rather show reactions similar to the period before the anticipation (isolation). In addition to confirm the ability in piglets to anticipate social reward, those behavioural data confirm the preference in isolated piglets for their conspecifics that represent a stronger positive valence than the arrival of a familiar human. Vocal expression differed between partners and were in line with behavioural observations. The inter-grunt interval was lower when piglets were expecting conspecifics. Morton’s rules explain that the rhythm of a behaviour can be positively linked to motivation^29^. Thus an increase in vocal activity when expecting conspecifics may be explained by the expression of a higher motivation toward this reward compared to the human reward, and thus a higher arousal.

This allowed us to measure the vocal expression of anticipation according to the event that is anticipated: arrival of pen mates or a familiar human (high familiarity after positive interactions versus lower familiarity). Considering the spectro-temporal features of grunts, although we failed in measuring a change in grunt duration during the anticipation of pen mates, we found an increase in duration after the signal stopped, when the entrance of pen mates was delayed. This latter phase is certainly a context with a negative valence (social isolation) and piglets may express the negative valence of this context in the duration of their grunt. In fact, several examples of the literature show that grunt duration is higher in negative contexts^18,28^. The duration of piglets’ grunt was not changed during anticipation. Two non-exclusive hypotheses can be raised to explain this result: 1) piglets learned that something positive was going to happen when entering the room (either the arrival of its penmates or the arrival of a human with positive contacts, but not an isolation) and expressed a non-specific positive state during the initial phase, so grunt produced during the initial phase are already ‘positive grunt’, 2) anatomical constraints of piglets’ vocal tract does not allow to shorten the grunts.

In a recent study, Briefer *et al*.^19^ showed that vocalisations (but not specifically grunts although they are usually over represented in datasets) recorded in positive contexts lasted 0.34 <0.42< 0.51 seconds. Considering all grunts produced during the initial phase in our study, the grunt duration is on average of 0.13 < 0.27 < 0.41 seconds (table S4). This may be in line with the first hypothesis, although the second hypothesis cannot be ruled out. To disentangle these hypotheses, we would need to measure grunt duration in a two-way associative learning with a positive and a negative social context (isolation vs. arrival of conspecifics), however the aim of the present study was to compare various positive contexts.

Piglets expecting a human already produced longer grunts during the anticipation phase, grunt that remained longer after the signal stopped, when the entrance of the human was delayed (similarly to when the entrance of penmates was delayed). An increase in grunt duration during the anticipation phase may mean that having a human as reward, instead of conspecifics may be perceive negatively by piglets. We found an average duration for the phase of (0.031 < 0.35 < 0.39, table S4), value that are similar to what Briefer *et al.*^19^ found for negative contexts. This result is surprising because our behavioural data showed that additional contacts had a positive effect on human-piglet relationship: piglets remained closer to the human (fig. S1). So for at least half of the piglets we can conclude that the presence of the human was positive compared to being isolated before the conditioning took place. In addition, the treatment had no effect on grunt duration comparing the initial and anticipation phases. So we can conclude a smoothing of the differences in familiarity degrees along the conditioning. Since anticipation extrapolate emotional states^30^ and that during the conditioning, two different output may arise in one trial (either human or conspecifics), we can hypothesize, that piglets may experience a cognitive bias. Piglets may rank the two possible outputs, increase the positivity of having conspecifics and increase the negativity of having a human arriving. In that case, vocal expression of anticipation of a human may be already the expression of a frustration rather than a positive emotion of lower intensity compared to anticipating conspecifics.

Spectral features of grunt changed drastically regarding the quality of the partner and contrary to grunt duration, changes were specific from the anticipation phase. Indeed, the spectral score increased when anticipating pen mates and decreased when anticipating the human but it returned to similar values as during the initial phase after the signal stopped, when the entrance of the partner was delayed. Therefore, the acoustic spectral score did not vary the same way depending on the quality of the partner: when expecting conspecifics, piglets produced grunt with higher spectral noise whereas they produced grunts with a higher frequency range and temporal noise when excepting a human. We can hypothesize that these rapid spectral changes are linked to rapid changes in the emotional arousal of piglets. Indeed harmonicity decreases with arousal in grunts^17^. Another measure would confirm such a hypothesis: for example heart-rate and its variability would be good indicators the arousal of the animals during anticipation phase and not for valence^12^.

To conclude, we showed that piglets were able to express behavioural and vocal flexibility when anticipating (pseudo)social events. In addition, grunts were spectrally and temporally different whether they were expecting a social reunion or an arrival of a familiar human. More interestingly, we also showed that analysing spectro-temporal properties of grunts allowed distinguishing between contexts (violation of expectations, positive human handling). Thus, acoustic analyses and especially of grunts, that are the most expressed type of vocalisations in pigs, allow tracking subtle changes in emotional states that behavioural analyses could not. Our original results in vocal features claim the possibly to better explore emotional states in non-verbal animals than a mere behavioural investigation.

## Authors contributions

Conceived and designed the experiment (A.V., C.T., A.B., C.G.). Performed the experiment (A.V., C.G., A.H., M.D.). Collection and edition of the acoustic and behavioural data (A.V., C.G., A.H., M.D.). Statistical analyses (A.V.). Contributed to the writing of the manuscript (A.V., C.T., A.B.).

## Acknowledgments

We acknowledge all the technical staff at UEPR: especially Patrick Touanel and Marie-Hélène Lohat, who largely participated in handling the piglets. We thank Eric Siroux who helped building the acoustic chamber at the beginning of the experiment and Remi Resmond for great discussion about statistics. This project is part of the SoundWel project in the framework of the Anihwa Eranet and funded by ANR 30001199.

## Data availability

The datasets used and/or analyzed during the current study are available from the corresponding author on reasonable request.

