## supplementary file for "Piglets vocally express the anticipation of (pseudo)-social contexts in their grunts"

### Title

### Supplementary data file

In this file are gathered all supplementary information not required to understand the study but provide more information than the main document and may be useful to replicate the experiment or use it for further meta analyses.

### Methods

ADDITIONAL HUMAN CONTACTS PROCEDURE

Similarly to Tallet *et al.*[1], the last phase was adapted to the reaction of each animal and included four steps: (1), the handler hold out the hand towards the animal; (2) if the piglet did not move away, the handler tried to touch it; (3) if the piglet accepted being touched, the handler softly stroked it along the body with the palm of her hand; and (4) once it accepted being stroked, the handler scratched it along the body with her fingers. Scratching consisted in rubbing the skin of the piglets with the finger tips and applying more pressure than stroking. Both experimenters trained, prior to the experiment, with non-experimental piglets to standardise the way they handled the piglets. In addition, the handler spoke to the piglet with a soft voice from the time she sat on the bucket until the end.

HUMAN-PIGLET REUNION STANDARD TRIAL BEFORE CONDITIONING

To trial the success of the additional human contact treatment to generate two groups of piglets having different degrees of familiarity toward the human, a standard trial of five-minute reunion with the human after five minutes of social isolation[2] was performed after the taming and before the conditioning.

### Results

ADDITIONAL HUMAN CONTACTS INCREASE HUMAN-PIGLET FAMILIARITY.

An Anova on the model testing the interaction between the treatment and the replicate, in addition to the experimenter identity showed no interaction between the treatment and the replicate ($X^2_1 = 0.005$, $p = 0.94$), a main effect of the treatment ($X^2_1 = 15.2$, $p = 0.0028$) and no effect of the experimenter ($X^2_1 = 0.51$, $p = 0.48$). Piglets that had received additional human contacts spent more time near the human (< 0.7 meter) (mean ± se, H+: 99.3 ± 12.4, H: 43.4 ± 11.9 seconds) over the five minutes. This trial allowed us to validate that H and H+ groups add different degrees of familiarity with the human (see additional figure S1) and to remove the factor of the experimenter in further analyses. This is a rapid validation of the taming protocol that have already been published before[2].

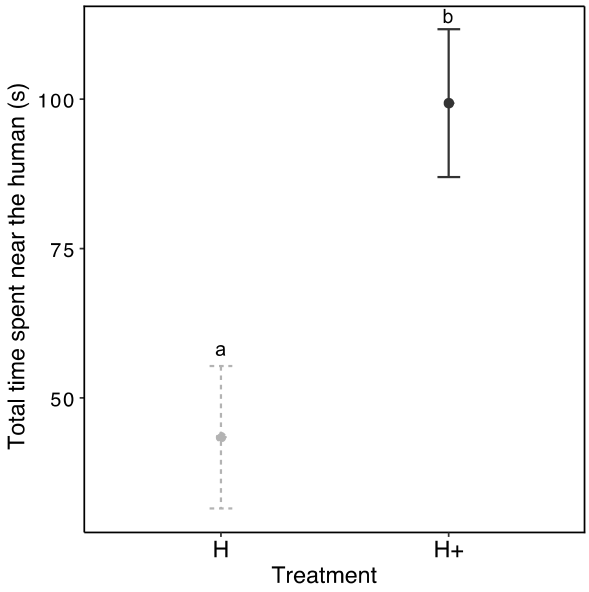

**Figure S1: Effect of post weaning additional human contact on piglet's proximity to a human**. Mean ± se per group (H: control, H+: additional human contact).

*Vocal types produced differs depending on the type of partner*

General vocal activity was studied using the number of calls of the different categories pigs do produce (grunts, squeals, screams, barks, mixed-calls) and inter call intervals. First, an analysis of the quality and quantity of call types depending on the partner, the phase of the trial and the treatment was carried out. Due to non-converging models (explained by the unbalanced occurrence of call types and the number of individuals contributing to them), no reliable statistics could be run. However, it has to be noticed that some call types only occurred in some groups: for instance, screams or squeals were exclusively associated to the human partner, bark were associated to the anticipation phase (phase 0, figure 2). This is the reason why, only grunts were considered in the study, since they were present in all groups and produced by the high enough number of individuals in each of them. This reflects the recent studies on vocal expression of identity or of emotions focusing on grunts [3-5]. Statistics showed no significant interactions between the treatment, the partner and the phase of the trial on the number of grunts produced but a significant effect of the phase (see table S1, S3). No effect of the partner nor treatment were found ($X^2_4=2.78$ $p=0.10$, $X^2_4=0.180$ $p=0.67$ respectively).

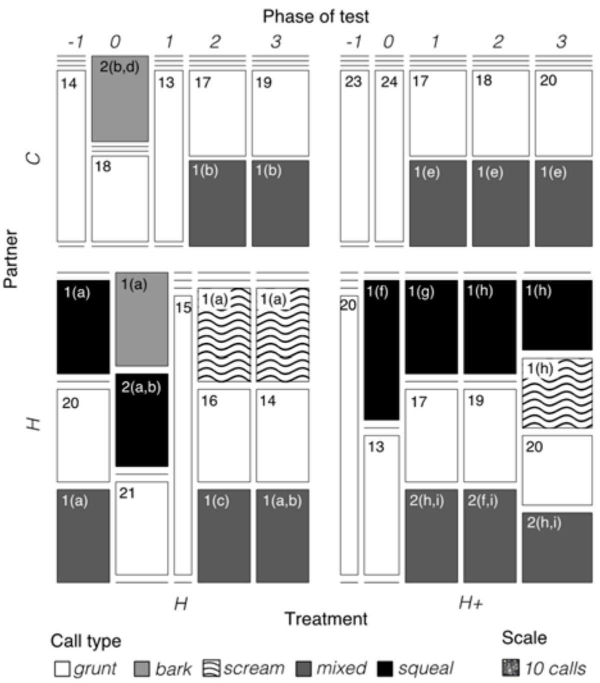

**Figure S2: General vocal activity during the anticipation trial according to the phase of the trial** (-1, 0, 1, 2 and 3)**, the type of partner** (C: conspecifics, H: human) **and the treatment** (H+: additional contacts, H: minimal contact). Phase -1 corresponds to the time before the broadcasting of a signal, phase 0 corresponds to the time during the signal and phases 1, 2, and 3 are 30 second segments during the violation of expectation period (90 seconds in total). Vocal activity was computed according to the number of calls of each call type produced per group and the rhythm of production. A: mosaic plot representing the relative proportion of each call type (grunt, squeal, scream, mixed, bark) according to the partner, the treatment and the phase. Number of each box represent the number of individuals contributing to the counting. When less than three individuals contributed, the identity of the piglet was also written in the box (a to i letters) to allow an illustration of repeated individuals for several categories. The scale of the number of calls illustrated by a square is also indicated. No statistics could be run to analyse the quality and quantity of calls types depending the partner and treatment due to non-converging models, but the number of grunts was.

STATISTICAL TABLES

**Table S1: Anova table for all models computed in the study**. After model validation, effects of explanatory variables were computed using the ‘Anova’ function (‘car’ R package), running Type II Wald chisquare trials. P values are labelled were significant (***: p<0.001, **: p<0.01, *: p<0.05, •:p<0.10). All linear models were computed using the ’lmer’ function, taking into account repeated observations as random factors. Models on counting were computed with a generalized model taking into account repeated observations as random factors, using a Poisson distribution (see model 4).

| | Chisq | Df | P value |
|---|---|---|---|
| ***model 1 : Behavioural Score 1 (LD1)*** | | | |
| Partner | 790.937 | 1 | <0.001*** |
| Phase of trial | 90.094 | 4 | <0.001*** |
| Treatment | 0.133 | 1 | 0.715 |
| Batch | 2.706 | 1 | 0.100 |
| Partner: Phase of trial | 13.929 | 4 | 0.008** |
| Partner : Treatment | 1.995 | 1 | 0.158 |
| Phase of trial : treatment | 0.688 | 4 | 0.953 |

| | | | |
|---|---|---|---|
| Partner : Batch | 0.035 | 1 | 0.852 |
| Treatment : Batch | 0.030 | 1 | 0.863 |
| Partner : Phase of trial : treatment | 5.232 | 4 | 0.264 |
| ***model 2 : Behavioural Score 2 (LD2)*** | | | |
| Partner | 5.590 | 1 | 0.018* |
| Phase of trial | 105.357 | 4 | <0.001*** |
| Treatment | 0.256 | 1 | 0.613 |
| Batch | 2.556 | 1 | 0.110 |
| Partner: Phase of trial | 49.066 | 4 | <0.001*** |
| Partner : Treatment | 0.069 | 1 | 0.793 |
| Phase of trial : Treatment | 2.594 | 4 | 0.628 |
| Partner : Batch | 0.301 | 1 | 0.583 |
| Treatment : Batch | 0.184 | 1 | 0.668 |
| Partner : Phase of trial : treatment | 1.824 | 4 | 0.768 |
| ***model 3 : log(Total time spent near upcoming partner's door +1)*** | | | |
| Trial (3 levels of factor) | 15.248 | 2 | <0.001*** |
| Partner | 252.790 | 1 | <0.001*** |
| Treatment | 3.789 | 1 | 0.052• |
| Batch | 0.373 | 1 | 0.542 |
| Trial: Partner | 11.956 | 2 | 0.003** |
| Trial: Treatment | 2.799 | 2 | 0.247 |
| Partner: Treatment | 0.000 | 1 | 0.999 |
| Partner: Batch | 0.227 | 1 | 0.634 |
| Treatment: Batch | 4.134 | 1 | 0.042* |
| Trial: Partner: Treatment | 5.208 | 2 | 0.074• |
| ***Model 4: Mean Inter Grunt Interval (log)*** | | | |
| Partner | 4.345 | 1 | 0.037* |
| Treatment | 1.519 | 1 | 0.218 |
| Batch | 0.113 | 1 | 0.737 |
| Partner : Treatment | 0.037 | 1 | 0.847 |
| Partner: Batch | 0.610 | 1 | 0.435 |
| Treatment : Batch | 0.155 | 1 | 0.693 |
| ***model 5 : Spectral acoustic Score (LD1)*** | | | |
| Partner | 22.778 | 1 | <0.001*** |
| Treatment | 3.527 | 1 | 0.060• |
| Phase of trial | 4.837 | 4 | 0.304 |
| Batch | 0.033 | 1 | 0.856 |
| Partner : Treatment | 6.453 | 1 | 0.01* |
| Partner: Phase of trial | 63.549 | 4 | <0.001*** |
| Phase of trial : Treatment | 14.762 | 4 | 0.005** |
| Treatment : Batch | 2.305 | 1 | 0.129 |
| Partner : Batch | 12.923 | 1 | <0.001*** |
| Partner : Phase of trial : treatment | 4.207 | 4 | 0.379 |
| ***model 6 : log(Vocalisation duration)*** | | | |
| Partner | 16.434 | 1 | <0.001*** |
| Treatment | 3.964 | 1 | 0.046* |
| Phase of trial | 65.777 | 4 | <0.001*** |
| Batch | 11.948 | 1 | 0.001** |
| Partner : Treatment | 5.843 | 1 | 0.016* |
| Partner: Phase of trial | 50.339 | 4 | <0.001*** |
| Phase of trial : Treatment | 12.414 | 4 | 0.015* |
| Treatment : Batch | 3.841 | 1 | 0.050• |
| Partner : Batch | 4.595 | 1 | 0.032* |

| | | | |
|---|---|---|---|
| Partner : Phase of trial : treatment | 6.327 | 4 | 0.176 |

**Table S1: Table of model estimates, following significant effect of explanatory variables**. All estimates were calculated using the 'lsmeans' function ('LmerTrial' R package) and are presented per relevant group lsmean, SE, DF, lower.CI upper.CI respectively represent mean estimates, standard error, degrees of freedom and lower and upper limit of 95% confidence interval.

| Variable 1 | Variable 2 | lsmean | SE | DF | Lower.CI | Upper.CI |
|---|---|---|---|---|---|---|
| ***model 1 : behavioural Score 1 (LD1)*** | | | | | | |
| Partner | Phase of trial | | | | | |
| C | -1 | -0.411 | 0.080 | 436.097 | -0.568 | -0.255 |
| H | -1 | 1.094 | 0.079 | 433.456 | 0.938 | 1.250 |
| C | 0 | -1.171 | 0.081 | 456.643 | -1.330 | -1.011 |
| H | 0 | 0.477 | 0.081 | 453.933 | 0.318 | 0.637 |
| C | 1 | -0.778 | 0.129 | 940.783 | -1.032 | -0.524 |
| H | 1 | 0.716 | 0.129 | 939.334 | 0.462 | 0.970 |
| C | 2 | -0.859 | 0.129 | 940.783 | -1.113 | -0.605 |
| H | 2 | 1.040 | 0.129 | 939.334 | 0.786 | 1.294 |
| C | 3 | -1.153 | 0.129 | 940.783 | -1.407 | -0.899 |
| H | 3 | 1.032 | 0.129 | 939.334 | 0.778 | 1.286 |
| ***model 2 : behavioural Score 2 (LD2)*** | | | | | | |
| Partner | Phase of trial | | | | | |
| C | -1 | -0.313 | 0.074 | 716.748 | -0.458 | -0.168 |
| H | -1 | -0.426 | 0.074 | 714.272 | -0.571 | -0.282 |
| C | 0 | -0.002 | 0.076 | 731.417 | -0.150 | 0.147 |
| H | 0 | 0.742 | 0.075 | 729.046 | 0.594 | 0.890 |
| C | 1 | -0.014 | 0.126 | 1016.193 | -0.262 | 0.233 |
| H | 1 | -0.185 | 0.126 | 1015.277 | -0.432 | 0.063 |
| C | 2 | 0.075 | 0.126 | 1016.193 | -0.172 | 0.323 |
| H | 2 | -0.118 | 0.126 | 1015.277 | -0.365 | 0.130 |
| *C* | 3 | 0.214 | 0.126 | 1016.193 | -0.033 | 0.461 |
| H | 3 | 0.036 | 0.126 | 1015.277 | -0.212 | 0.283 |
| ***model 3 : log( Total time spent near upcoming partner's door +1)*** | | | | | | |
| Trial | Partner | | | | | |
| Beginning | C | 1.123 | 0.094 | 204.997 | 0.939 | 1.308 |
| Middle | C | 1.115 | 0.106 | 433.606 | 0.906 | 1.324 |
| End | C | 1.693 | 0.081 | 161.538 | 1.534 | 1.852 |
| Beginning | H | 0.210 | 0.092 | 194.679 | 0.030 | 0.391 |
| Middle | H | 0.243 | 0.106 | 436.376 | 0.034 | 0.453 |
| End | H | 0.309 | 0.081 | 160.583 | 0.151 | 0.468 |
| Treatment | Batch | | | | | |
| H | 1 | 0.799 | 0.082 | 152.819 | 0.638 | 0.960 |
| H+ | 1 | 0.814 | 0.078 | 116.846 | 0.659 | 0.968 |
| H | 2 | 0.907 | 0.069 | 106.638 | 0.771 | 1.042 |
| H+ | 2 | 0.618 | 0.070 | 109.605 | 0.479 | 0.757 |
| ***model 4 : Mean Inter Grunt Interval (log)*** | | | | | | |
| Partner | | | | | | |
| C | - | 1.231524626 | 0.061349464 | 66.37954653 | 1.109049541 | 1.353999711 |
| H | - | 1.368648332 | 0.059002793 | 64.47708243 | 1.250858026 | 1.486438637 |
| ***model 6 : Spectral acoustic Score 1 (LD1)*** | | | | | | |
| Partner | Phase of trial | | | | | |

| | | | | | | |
|---|---|---|---|---|---|---|
| C | -1 | 0.102 | 0.104 | 285.030 | -0.101 | 0.306 |
| H | -1 | 0.091 | 0.103 | 287.347 | -0.112 | 0.295 |
| C | 0 | 0.510 | 0.091 | 172.830 | 0.332 | 0.688 |
| H | 0 | -0.322 | 0.091 | 180.161 | -0.502 | -0.142 |
| C | 1 | 0.038 | 0.083 | 124.813 | -0.126 | 0.202 |
| H | 1 | -0.017 | 0.078 | 99.422 | -0.171 | 0.137 |
| C | 2 | 0.045 | 0.093 | 193.451 | -0.138 | 0.229 |
| H | 2 | -0.047 | 0.094 | 198.219 | -0.232 | 0.137 |
| C | 3 | 0.043 | 0.089 | 161.491 | -0.132 | 0.218 |
| H | 3 | 0.127 | 0.091 | 179.062 | -0.052 | 0.305 |
| Partner | Batch | | | | | |
| C | 1 | 0.249 | 0.096 | 60.346 | 0.058 | 0.440 |
| H | 1 | -0.092 | 0.094 | 55.780 | -0.280 | 0.095 |
| C | 2 | 0.047 | 0.102 | 66.153 | -0.158 | 0.251 |
| H | 2 | 0.025 | 0.102 | 68.424 | -0.180 | 0.229 |
| Partner | Treatment | | | | | |
| C | H | 0.058 | 0.103 | 70.509 | -0.147 | 0.263 |
| H | H | -0.223 | 0.102 | 70.788 | -0.426 | -0.021 |
| C | H+ | 0.237 | 0.096 | 57.934 | 0.046 | 0.429 |
| H | H+ | 0.156 | 0.095 | 56.403 | -0.035 | 0.346 |
| Treatment | Phase of trial | | | | | |
| H | -1 | -0.166 | 0.127 | 164.945 | -0.417 | 0.085 |
| H+ | -1 | 0.359 | 0.116 | 120.145 | 0.131 | 0.588 |
| H | 0 | 0.037 | 0.118 | 122.512 | -0.195 | 0.270 |
| H+ | 0 | 0.151 | 0.103 | 77.191 | -0.053 | 0.355 |
| H | 1 | -0.045 | 0.104 | 76.524 | -0.251 | 0.161 |
| H+ | 1 | 0.066 | 0.100 | 67.989 | -0.131 | 0.264 |
| H | 2 | -0.118 | 0.115 | 111.674 | -0.344 | 0.108 |
| H+ | 2 | 0.116 | 0.108 | 94.744 | -0.098 | 0.330 |
| H | 3 | -0.120 | 0.112 | 103.783 | -0.341 | 0.100 |
| H+ | 3 | 0.290 | 0.104 | 81.848 | 0.084 | 0.496 |
| ***model 7 : log( Vocalisation duration)*** | | | | | | |
| Partner | Phase of trial | | | | | |
| C | -1 | -1.398 | 0.055 | 243.478 | -1.506 | -1.290 |
| H | -1 | -1.390 | 0.055 | 244.706 | -1.497 | -1.282 |
| C | 0 | -1.525 | 0.048 | 152.502 | -1.620 | -1.430 |
| H | 0 | -1.135 | 0.049 | 158.111 | -1.231 | -1.039 |
| C | 1 | -1.215 | 0.045 | 113.647 | -1.303 | -1.126 |
| H | 1 | -1.204 | 0.042 | 92.526 | -1.287 | -1.120 |
| C | 2 | -1.134 | 0.050 | 169.262 | -1.231 | -1.036 |
| H | 2 | -1.117 | 0.050 | 172.929 | -1.215 | -1.019 |
| C | 3 | -1.186 | 0.047 | 143.129 | -1.279 | -1.092 |
| H | 3 | -1.195 | 0.048 | 157.323 | -1.290 | -1.100 |
| Partner | Batch | | | | | |
| C | 1 | -1.446 | 0.053 | 59.658 | -1.552 | -1.341 |
| H | 1 | -1.314 | 0.052 | 55.732 | -1.418 | -1.211 |
| C | 2 | -1.137 | 0.056 | 65.088 | -1.249 | -1.024 |
| H | 2 | -1.102 | 0.056 | 66.706 | -1.214 | -0.990 |
| Partner | Treatment | | | | | |
| C | H | -1.241 | 0.056 | 69.082 | -1.353 | -1.129 |
| H | H | -1.105 | 0.056 | 68.812 | -1.216 | -0.994 |
| C | H+ | -1.342 | 0.053 | 57.403 | -1.448 | -1.237 |
| H | H+ | -1.311 | 0.053 | 56.159 | -1.416 | -1.206 |
| Treatment | Batch | | | | | |

| | | | | | | |
|---|---|---|---|---|---|---|
| H | 1 | -1.375 | 0.068 | 46.746 | -1.512 | -1.237 |
| H+ | 1 | -1.386 | 0.073 | 50.125 | -1.532 | -1.239 |
| H | 2 | -0.971 | 0.081 | 63.407 | -1.133 | -0.809 |
| H+ | 2 | -1.268 | 0.070 | 45.241 | -1.408 | -1.127 |
| Treatment | Phase of trial | | | | | |
| H | -1 | -1.298 | 0.068 | 147.175 | -1.433 | -1.164 |
| H+ | -1 | -1.490 | 0.062 | 108.741 | -1.613 | -1.366 |
| H | 0 | -1.311 | 0.063 | 112.239 | -1.437 | -1.186 |
| H+ | 0 | -1.348 | 0.056 | 73.494 | -1.459 | -1.237 |
| H | 1 | -1.157 | 0.057 | 73.946 | -1.269 | -1.044 |
| H+ | 1 | -1.261 | 0.055 | 65.900 | -1.370 | -1.153 |
| H | 2 | -1.021 | 0.062 | 102.837 | -1.143 | -0.899 |
| H+ | 2 | -1.230 | 0.059 | 87.948 | -1.346 | -1.113 |
| H | 3 | -1.077 | 0.060 | 95.992 | -1.196 | -0.957 |
| H+ | 3 | -1.304 | 0.057 | 77.183 | -1.417 | -1.192 |

**Table S2: Post hoc trials on models following significant interactions or single effect of one or more explanatory variable(s)**. Eah model is indicated, all post hoc trials were computed using the ’lsmeans’ function with Tukey correction for multiple trialing. All relevant contrasts were bolded to increase visibility and significant p values indicated (***: p<0.001, **: p<0.01, *: p<0.05, • :p<0.10).

| Contrast | Estimate | SE | DF | T.ratio | P.value |
|---|---|---|---|---|---|
| ***model 1 : behavioural Score 1 (LD1)*** | | | | | |
| Partner: Phase of trial interaction | | | | | |
| **C,-1 - H,-1** | **-1.505** | **0.102** | **966.348** | **-14.742** | **<0.001******* |
| **C,-1 - C,0** | **0.759** | **0.104** | **968.786** | **7.331** | **<0.001******* |
| C,-1 - H,0 | -0.889 | 0.103 | 968.908 | -8.594 | <0.001*** |
| **C,-1 - C,1** | **0.367** | **0.144** | **966.275** | **2.543** | **0.247** |
| C,-1 - H,1 | -1.127 | 0.144 | 967.465 | -7.809 | <0.001*** |
| **C,-1 - C,2** | **0.448** | **0.144** | **966.275** | **3.104** | **0.061•** |
| C,-1 - H,2 | -1.451 | 0.144 | 967.465 | -10.053 | <0.001*** |
| **C,-1 - C,3** | **0.742** | **0.144** | **966.275** | **5.140** | **<0.001******* |
| C,-1 - H,3 | -1.444 | 0.144 | 967.465 | -9.999 | <0.001*** |
| H,-1 - C,0 | 2.265 | 0.103 | 968.923 | 21.899 | <0.001*** |
| **H,-1 - H,0** | **0.617** | **0.103** | **968.767** | **5.973** | **<0.001******* |
| H,-1 - C,1 | 1.872 | 0.144 | 966.205 | 12.984 | <0.001*** |
| **H,-1 - H,1** | **0.378** | **0.144** | **967.402** | **2.621** | **0.209** |
| H,-1 - C,2 | 1.953 | 0.144 | 966.205 | 13.545 | <0.001*** |
| **H,-1 - H,2** | **0.054** | **0.144** | **967.402** | **0.375** | **1.000** |
| H,-1 - C,3 | 2.247 | 0.144 | 966.205 | 15.583 | <0.001*** |
| **H,-1 - H,3** | **0.062** | **0.144** | **967.402** | **0.429** | **1.000** |
| **C,0 - H,0** | **-1.648** | **0.105** | **966.346** | **-15.765** | **<0.001******* |
| **C,0 - C,1** | **-0.392** | **0.145** | **967.590** | **-2.701** | **0.175** |
| C,0 - H,1 | -1.886 | 0.145 | 968.907 | -12.984 | <0.001*** |
| **C,0 - C,2** | **-0.311** | **0.145** | **967.590** | **-2.143** | **0.497** |
| C,0 - H,2 | -2.210 | 0.145 | 968.907 | -15.215 | <0.001*** |
| **C,0 - C,3** | **-0.017** | **0.145** | **967.590** | **-0.120** | **1.000** |
| C,0 - H,3 | -2.203 | 0.145 | 968.907 | -15.161 | <0.001*** |
| H,0 - C,1 | 1.256 | 0.145 | 967.509 | 8.652 | <0.001*** |
| **H,0 - H,1** | **-0.239** | **0.145** | **968.824** | **-1.644** | **0.826** |
| H,0 - C,2 | 1.336 | 0.145 | 967.509 | 9.210 | <0.001*** |

| Contrast | Estimate | SE | DF | T.ratio | P.value |
|---|---|---|---|---|---|
| ***model 2 : behavioural Score 2 (LD2)*** | | | | | |
| Partner: Phase of trial interaction | | | | | |
| **C,-1 - H,-1** | **0.113** | **0.102** | **967.064** | **1.105** | **0.984** |
| **C,-1 - C,0** | **-0.312** | **0.104** | **970.812** | **-3.005** | **0.081•** |
| C,-1 - H,0 | -1.055 | 0.104 | 971.017 | -10.187 | <0.001*** |
| **C,-1 - C,1** | **-0.299** | **0.145** | **966.940** | **-2.068** | **0.550** |
| C,-1 - H,1 | -0.128 | 0.145 | 969.040 | -0.887 | 0.997 |
| **C,-1 - C,2** | **-0.389** | **0.145** | **966.940** | **-2.687** | **0.180** |
| C,-1 - H,2 | -0.195 | 0.145 | 969.040 | -1.350 | 0.942 |
| **C,-1 - C,3** | **-0.527** | **0.145** | **966.940** | **-3.645** | **0.010*** |
| C,-1 - H,3 | -0.349 | 0.145 | 969.040 | -2.414 | 0.319 |
| H,-1 - C,0 | -0.425 | 0.104 | 971.041 | -4.101 | 0.002** |
| **H,-1 - H,0** | **-1.168** | **0.103** | **970.782** | **-11.294** | **<0.001******* |
| H,-1 - C,1 | -0.412 | 0.145 | 966.822 | -2.853 | 0.121 |
| **H,-1 - H,1** | **-0.242** | **0.145** | **968.932** | **-1.671** | **0.812** |
| H,-1 - C,2 | -0.502 | 0.145 | 966.822 | -3.472 | 0.019* |
| **H,-1 - H,2** | **-0.308** | **0.145** | **968.932** | **-2.134** | **0.504** |
| H,-1 - C,3 | -0.640 | 0.145 | 966.822 | -4.431 | <0.001*** |
| **H,-1 - H,3** | **-0.462** | **0.145** | **968.932** | **-3.198** | **0.046*** |
| **C,0 - H,0** | **-0.744** | **0.105** | **967.060** | **-7.097** | **<0.001******* |
| **C,0 - C,1** | **0.013** | **0.146** | **968.976** | **0.086** | **1.000** |
| C,0 - H,1 | 0.183 | 0.146 | 971.225 | 1.260 | 0.962 |
| **C,0 - C,2** | **-0.077** | **0.146** | **968.976** | **-0.529** | **1.000** |
| C,0 - H,2 | 0.116 | 0.146 | 971.225 | 0.800 | 0.999 |
| **C,0 - C,3** | **-0.216** | **0.146** | **968.976** | **-1.481** | **0.900** |
| C,0 - H,3 | -0.037 | 0.146 | 971.225 | -0.257 | 1.000 |
| H,0 - C,1 | 0.756 | 0.145 | 968.843 | 5.199 | <0.001*** |
| **H,0 - H,1** | **0.927** | **0.145** | **971.091** | **6.373** | **<0.001******* |
| H,0 - C,2 | 0.667 | 0.145 | 968.843 | 4.584 | <0.001*** |

| | | | | | |
|---|---|---|---|---|---|
| H,0 - H,2 | -0.563 | 0.145 | 968.824 | -3.876 | 0.004** |
| H,0 - C,3 | 1.630 | 0.145 | 967.509 | 11.235 | <0.001*** |
| H,0 - H,3 | -0.555 | 0.145 | 968.824 | -3.823 | 0.005** |
| C,1 - H,1 | -1.494 | 0.177 | 967.001 | -8.458 | <0.001*** |
| C,1 - C,2 | 0.081 | 0.177 | 966.205 | 0.458 | 1.000 |
| C,1 - H,2 | -1.818 | 0.177 | 967.001 | -10.292 | <0.001*** |
| C,1 - C,3 | 0.375 | 0.177 | 966.205 | 2.122 | 0.512 |
| C,1 - H,3 | -1.810 | 0.177 | 967.001 | -10.248 | <0.001*** |
| H,1 - C,2 | 1.575 | 0.177 | 967.001 | 8.916 | <0.001*** |
| H,1 - H,2 | -0.324 | 0.177 | 966.205 | -1.835 | 0.713 |
| H,1 - C,3 | 1.869 | 0.177 | 967.001 | 10.580 | <0.001*** |
| H,1 - H,3 | -0.316 | 0.177 | 966.205 | -1.791 | 0.741 |
| C,2 - H,2 | -1.899 | 0.177 | 967.001 | -10.751 | <0.001*** |
| C,2 - C,3 | 0.294 | 0.177 | 966.205 | 1.664 | 0.816 |
| C,2 - H,3 | -1.891 | 0.177 | 967.001 | -10.707 | <0.001*** |
| H,2 - C,3 | 2.193 | 0.177 | 967.001 | 12.414 | <0.001*** |
| H,2 - H,3 | 0.008 | 0.177 | 966.205 | 0.044 | 1.000 |
| C,3 - H,3 | -2.185 | 0.177 | 967.001 | -12.370 | <0.001*** |

***model 3 : log( Total time spent near upcoming partner's door +1)***

Partner: Trial (3 levels of factor)

| | | | | | |
|---|---|---|---|---|---|
| beginning,C - end,C | -0.570 | 0.125 | 189.673 | -4.546 | <0.001*** |
| beginning,C - middle,C | 0.008 | 0.140 | 527.352 | 0.057 | 1.000 |
| beginning,C - beginning,H | 0.913 | 0.130 | 639.411 | 7.021 | <0.001*** |
| beginning,C - end,H | 0.814 | 0.125 | 189.230 | 6.498 | <0.001*** |
| beginning,C - middle,H | 0.880 | 0.141 | 528.253 | 6.250 | <0.001*** |
| end,C - middle,C | 0.578 | 0.138 | 197.543 | 4.187 | 0.001** |
| end,C - beginning,H | 1.483 | 0.124 | 182.762 | 11.980 | <0.001*** |
| end,C - end,H | 1.384 | 0.103 | 615.691 | 13.438 | <0.001*** |
| end,C - middle,H | 1.450 | 0.138 | 197.204 | 10.512 | <0.001*** |
| middle,C - beginning,H | 0.905 | 0.139 | 524.240 | 6.506 | <0.001*** |
| middle,C - end,H | 0.806 | 0.137 | 195.104 | 5.862 | <0.001*** |
| middle,C - middle,H | 0.872 | 0.148 | 616.184 | 5.901 | <0.001*** |
| beginning,H - end,H | -0.099 | 0.124 | 181.829 | -0.802 | 0.967 |
| beginning,H - middle,H | -0.033 | 0.140 | 527.302 | -0.236 | 1.000 |
| end,H - middle,H | 0.066 | 0.138 | 199.068 | 0.478 | 0.997 |

| | | | | | |
|---|---|---|---|---|---|
| H,0 - H,2 | 0.860 | 0.145 | 971.091 | 5.913 | <0.001*** |
| H,0 - C,3 | 0.528 | 0.145 | 968.843 | 3.631 | 0.011* |
| H,0 - H,3 | 0.706 | 0.145 | 971.091 | 4.855 | <0.001*** |
| C,1 - H,1 | 0.171 | 0.177 | 968.229 | 0.965 | 0.994 |
| C,1 - C,2 | -0.090 | 0.177 | 966.822 | -0.506 | 1.000 |
| C,1 - H,2 | 0.104 | 0.177 | 968.229 | 0.587 | 1.000 |
| C,1 - C,3 | -0.228 | 0.177 | 966.822 | -1.288 | 0.956 |
| C,1 - H,3 | -0.050 | 0.177 | 968.229 | -0.282 | 1.000 |
| H,1 - C,2 | -0.260 | 0.177 | 968.229 | -1.470 | 0.904 |
| H,1 - H,2 | -0.067 | 0.177 | 966.822 | -0.378 | 1.000 |
| H,1 - C,3 | -0.399 | 0.177 | 968.229 | -2.253 | 0.421 |
| H,1 - H,3 | -0.221 | 0.177 | 966.822 | -1.247 | 0.964 |
| C,2 - H,2 | 0.193 | 0.177 | 968.229 | 1.092 | 0.985 |
| C,2 - C,3 | -0.139 | 0.177 | 966.822 | -0.783 | 0.999 |
| C,2 - H,3 | 0.040 | 0.177 | 968.229 | 0.223 | 1.000 |
| H,2 - C,3 | -0.332 | 0.177 | 968.229 | -1.875 | 0.686 |
| H,2 - H,3 | -0.154 | 0.177 | 966.822 | -0.869 | 0.997 |
| C,3 - H,3 | 0.178 | 0.177 | 968.229 | 1.006 | 0.992 |

***model 3 : log( Total time spent near upcoming partner's door +1)***

Treatment: Batch interaction

| | | | | | |
|---|---|---|---|---|---|
| H,1 - H+,1 | -0.014 | 0.113 | 134.234 | -0.128 | 0.999 |
| H,1 - H,2 | -0.107 | 0.106 | 108.450 | -1.014 | 0.742 |
| H,1 - H+,2 | 0.181 | 0.108 | 133.567 | 1.677 | 0.340 |
| H+,1 - H,2 | -0.093 | 0.104 | 113.122 | -0.893 | 0.808 |
| H+,1 - H+,2 | 0.195 | 0.105 | 91.903 | 1.866 | 0.250 |
| H,2 - H+,2 | 0.288 | 0.098 | 108.150 | 2.932 | 0.021* |

***model 5 : Spectral acoustic Score (LD1)***

Partner: Phase of trial interaction

| | | | | | |
|---|---|---|---|---|---|
| **C,-1 - H,-1** | **0.011** | **0.115** | **2217.203** | **0.098** | **1.000** |
| **C,-1 - C,0** | **-0.408** | **0.103** | **2209.822** | **-3.965** | **0.003**** |
| C,-1 - H,0 | 0.424 | 0.105 | 2216.153 | 4.043 | 0.002** |
| **C,-1 - C,1** | **0.064** | **0.097** | **2208.760** | **0.664** | **1.000** |
| C,-1 - H,1 | 0.120 | 0.095 | 2229.751 | 1.257 | 0.963 |
| **C,-1 - C,2** | **0.057** | **0.106** | **2213.284** | **0.537** | **1.000** |
| C,-1 - H,2 | 0.150 | 0.109 | 2236.493 | 1.375 | 0.935 |
| **C,-1 - C,3** | **0.060** | **0.104** | **2222.636** | **0.576** | **1.000** |
| C,-1 - H,3 | -0.024 | 0.107 | 2241.714 | -0.227 | 1.000 |
| H,-1 - C,0 | -0.419 | 0.104 | 2217.079 | -4.023 | 0.002** |
| **H,-1 - H,0** | **0.413** | **0.105** | **2218.536** | **3.951** | **0.003**** |
| H,-1 - C,1 | 0.053 | 0.098 | 2219.878 | 0.540 | 1.000 |
| **H,-1 - H,1** | **0.108** | **0.094** | **2224.328** | **1.153** | **0.979** |
| H,-1 - C,2 | 0.046 | 0.108 | 2227.904 | 0.425 | 1.000 |
| **H,-1 - H,2** | **0.139** | **0.108** | **2230.613** | **1.287** | **0.957** |
| H,-1 - C,3 | 0.048 | 0.104 | 2230.920 | 0.463 | 1.000 |
| **H,-1 - H,3** | **-0.036** | **0.106** | **2239.583** | **-0.335** | **1.000** |
| **C,0 - H,0** | **0.832** | **0.092** | **2213.600** | **9.005** | **<0.001***** |
| **C,0 - C,1** | **0.472** | **0.083** | **2209.145** | **5.675** | **<0.001***** |
| C,0 - H,1 | 0.527 | 0.081 | 2234.726 | 6.502 | <0.001*** |
| **C,0 - C,2** | **0.465** | **0.095** | **2222.188** | **4.903** | **<0.001***** |
| C,0 - H,2 | 0.558 | 0.096 | 2236.855 | 5.782 | <0.001*** |
| **C,0 - C,3** | **0.467** | **0.091** | **2223.392** | **5.147** | **<0.001***** |
| C,0 - H,3 | 0.383 | 0.094 | 2244.946 | 4.064 | 0.002** |
| H,0 - C,1 | -0.360 | 0.086 | 2224.225 | -4.200 | 0.001** |
| **H,0 - H,1** | **-0.305** | **0.081** | **2231.749** | **-3.744** | **0.007**** |
| H,0 - C,2 | -0.367 | 0.096 | 2225.966 | -3.842 | 0.005** |
| **H,0 - H,2** | **-0.274** | **0.096** | **2231.261** | **-2.850** | **0.121** |
| H,0 - C,3 | -0.365 | 0.092 | 2228.687 | -3.967 | 0.003** |
| **H,0 - H,3** | **-0.449** | **0.095** | **2242.615** | **-4.723** | **<0.001***** |
| **C,1 - H,1** | **0.055** | **0.073** | **2240.836** | **0.761** | **0.999** |
| **C,1 - C,2** | **-0.007** | **0.087** | **2214.775** | **-0.083** | **1.000** |
| C,1 - H,2 | 0.086 | 0.090 | 2242.541 | 0.953 | 0.995 |

***model 5 : Spectral acoustic Score (LD1)***

Treatment: Phase of trial interaction

| | | | | | |
|---|---|---|---|---|---|
| **H,-1 - H+,-1** | **-0.525** | **0.172** | **142.068** | **-3.055** | **0.078•** |
| **H,-1 - H,0** | **-0.203** | **0.112** | **2215.443** | **-1.804** | **0.733** |
| H,-1 - H+,0 | -0.317 | 0.164 | 118.129 | -1.935 | 0.645 |
| **H,-1 - H,1** | **-0.121** | **0.101** | **2220.257** | **-1.200** | **0.973** |
| H,-1 - H+,1 | -0.232 | 0.162 | 112.282 | -1.434 | 0.914 |
| **H,-1 - H,2** | **-0.048** | **0.114** | **2232.977** | **-0.418** | **1.000** |
| **H,-1 - H+,2** | **-0.282** | **0.167** | **128.115** | **-1.687** | **0.800** |
| **H,-1 - H,3** | **-0.045** | **0.113** | **2237.855** | **-0.401** | **1.000** |
| H,-1 - H+,3 | -0.456 | 0.164 | 120.742 | -2.772 | 0.158 |
| H+,-1 - H,0 | 0.322 | 0.165 | 121.336 | 1.952 | 0.634 |
| **H+,-1 - H+,0** | **0.208** | **0.094** | **2214.486** | **2.207** | **0.452** |
| H+,-1 - H,1 | 0.404 | 0.156 | 96.944 | 2.597 | 0.233 |
| **H+,-1 - H+,1** | **0.293** | **0.090** | **2210.346** | **3.255** | **0.038*** |
| H+,-1 - H,2 | 0.478 | 0.163 | 115.836 | 2.934 | 0.108 |
| **H+,-1 - H+,2** | **0.243** | **0.100** | **2212.764** | **2.423** | **0.313** |
| H+,-1 - H,3 | 0.480 | 0.161 | 111.828 | 2.987 | 0.095• |
| **H+,-1 - H+,3** | **0.069** | **0.097** | **2216.499** | **0.714** | **0.999** |
| **H,0 - H+,0** | **-0.114** | **0.157** | **99.103** | **-0.729** | **0.999** |
| **H,0 - H,1** | **0.082** | **0.090** | **2226.312** | **0.917** | **0.996** |
| H,0 - H+,1 | -0.029 | 0.155 | 93.729 | -0.189 | 1.000 |
| **H,0 - H,2** | **0.155** | **0.103** | **2231.712** | **1.503** | **0.891** |
| H,0 - H+,2 | -0.079 | 0.160 | 108.272 | -0.494 | 1.000 |
| **H,0 - H,3** | **0.158** | **0.102** | **2238.722** | **1.540** | **0.876** |
| H,0 - H+,3 | -0.253 | 0.157 | 101.416 | -1.608 | 0.841 |
| H+,0 - H,1 | 0.196 | 0.147 | 76.927 | 1.338 | 0.941 |
| **H+,0 - H+,1** | **0.085** | **0.075** | **2223.071** | **1.134** | **0.981** |
| H+,0 - H,2 | 0.269 | 0.154 | 94.007 | 1.747 | 0.765 |
| **H+,0 - H+,2** | **0.035** | **0.086** | **2217.711** | **0.405** | **1.000** |
| H+,0 - H,3 | 0.272 | 0.152 | 90.068 | 1.788 | 0.741 |
| **H+,0 - H+,3** | **-0.139** | **0.082** | **2213.225** | **-1.695** | **0.799** |
| **H,1 - H+,1** | **-0.111** | **0.145** | **72.230** | **-0.770** | **0.999** |
| **H,1 - H,2** | **0.073** | **0.088** | **2224.569** | **0.835** | **0.998** |
| H,1 - H+,2 | -0.161 | 0.150 | 85.153 | -1.072 | 0.986 |

| | | | | | |
|---|---|---|---|---|---|
| **C,1 - C,3** | **-0.005** | **0.083** | **2224.133** | **-0.055** | **1.000** |
| C,1 - H,3 | -0.089 | 0.087 | 2246.887 | -1.016 | 0.991 |
| H,1 - C,2 | -0.063 | 0.085 | 2243.554 | -0.735 | 0.999 |
| **H,1 - H,2** | **0.030** | **0.082** | **2218.897** | **0.369** | **1.000** |
| H,1 - C,3 | -0.060 | 0.081 | 2246.462 | -0.737 | 0.999 |
| **H,1 - H,3** | **-0.144** | **0.080** | **2235.714** | **-1.789** | **0.742** |
| **C,2 - H,2** | **0.093** | **0.100** | **2243.084** | **0.928** | **0.996** |
| **C,2 - C,3** | **0.003** | **0.092** | **2210.153** | **0.029** | **1.000** |
| C,2 - H,3 | -0.081 | 0.098 | 2246.993 | -0.829 | 0.998 |
| H,2 - C,3 | -0.090 | 0.097 | 2245.410 | -0.933 | 0.995 |
| **H,2 - H,3** | **-0.174** | **0.094** | **2220.675** | **-1.853** | **0.701** |
| **C,3 - H,3** | **-0.084** | **0.095** | **2245.497** | **-0.888** | **0.997** |

Partner: Treatment interaction

| | | | | | |
|---|---|---|---|---|---|
| **C,H - H,H** | **0.281** | **0.070** | **2227.517** | **4.018** | **<0.001*** ** |
| **C,H - C,H+** | **-0.179** | **0.141** | **64.437** | **-1.273** | **0.583** |
| C,H - H,H+ | -0.098 | 0.140 | 63.322 | -0.696 | 0.898 |
| H,H - C,H+ | -0.461 | 0.139 | 63.808 | -3.303 | 0.008** |
| **H,H - H,H+** | **-0.379** | **0.140** | **63.571** | **-2.717** | **0.04*** |
| **C,H+ - H,H+** | **0.082** | **0.060** | **2245.888** | **1.369** | **0.519** |

***model 6 : log( Vocalisation duration)***

Partner: Phase of trial interaction

| | | | | | |
|---|---|---|---|---|---|
| **C,-1 - H,-1** | **-0.008** | **0.059** | **2214.509** | **-0.141** | **1.000** |
| **C,-1 - C,0** | **0.127** | **0.052** | **2207.760** | **2.418** | **0.315** |
| C,-1 - H,0 | -0.263 | 0.053 | 2213.425 | -4.928 | <0.001*** |
| **C,-1 - C,1** | **-0.183** | **0.049** | **2206.907** | **-3.723** | **0.008**** |
| C,-1 - H,1 | -0.195 | 0.048 | 2225.623 | -4.017 | 0.002 |
| **C,-1 - C,2** | **-0.264** | **0.054** | **2211.248** | **-4.884** | **<0.001*** ** |
| C,-1 - H,2 | -0.281 | 0.056 | 2232.014 | -5.069 | <0.001*** |
| **C,-1 - C,3** | **-0.212** | **0.053** | **2219.543** | **-4.022** | **0.002**** |
| C,-1 - H,3 | -0.203 | 0.054 | 2238.763 | -3.728 | 0.008** |
| H,-1 - C,0 | 0.135 | 0.053 | 2214.689 | 2.543 | 0.246 |
| **H,-1 - H,0** | **-0.255** | **0.053** | **2215.989** | **-4.793** | **<0.001*** ** |
| H,-1 - C,1 | -0.175 | 0.050 | 2217.356 | -3.512 | 0.016* |
| **H,-1 - H,1** | **-0.186** | **0.048** | **2220.846** | **-3.897** | **0.004**** |
| H,-1 - C,2 | -0.256 | 0.055 | 2224.663 | -4.675 | <0.001*** |

| | | | | | |
|---|---|---|---|---|---|
| **H,1 - H,3** | **0.075** | **0.086** | **2233.648** | **0.876** | **0.997** |
| H,1 - H+,3 | -0.335 | 0.148 | 78.997 | -2.272 | 0.419 |
| H+,1 - H,2 | 0.185 | 0.152 | 88.741 | 1.213 | 0.968 |
| **H+,1 - H+,2** | **-0.050** | **0.082** | **2211.875** | **-0.613** | **1.000** |
| H+,1 - H,3 | 0.187 | 0.150 | 85.030 | 1.246 | 0.962 |
| **H+,1 - H+,3** | **-0.224** | **0.077** | **2216.851** | **-2.897** | **0.107** |
| **H,2 - H+,2** | **-0.234** | **0.158** | **102.963** | **-1.487** | **0.894** |
| **H,2 - H,3** | **0.002** | **0.098** | **2216.547** | **0.023** | **1.000** |
| H,2 - H+,3 | -0.408 | 0.155 | 96.229 | -2.636 | 0.216 |
| H+,2 - H,3 | 0.237 | 0.156 | 99.241 | 1.521 | 0.880 |
| **H+,2 - H+,3** | **-0.174** | **0.088** | **2211.712** | **-1.974** | **0.617** |
| **H,3 - H+,3** | **-0.411** | **0.153** | **92.588** | **-2.688** | **0.194** |

Partner: Batch interaction

| | | | | | |
|---|---|---|---|---|---|
| **C,1 - H,1** | **0.341** | **0.060** | **2229.718** | **5.706** | **<0.001*** ** |
| **C,1 - C,2** | **0.202** | **0.140** | **62.744** | **1.448** | **0.475** |
| C,1 - H,2 | 0.224 | 0.140 | 64.074 | 1.604 | 0.384 |
| H,1 - C,2 | -0.139 | 0.139 | 61.102 | -1.001 | 0.750 |
| **H,1 - H,2** | **-0.117** | **0.138** | **61.049** | **-0.846** | **0.832** |
| **C,2 - H,2** | **0.022** | **0.067** | **2244.870** | **0.327** | **0.988** |

***model 6 : log( Vocalisation duration)***

Treatment: Phase of trial interaction

| | | | | | |
|---|---|---|---|---|---|
| **H,-1 - H+,-1** | **0.191** | **0.092** | **127.492** | **2.074** | **0.549** |
| **H,-1 - H,0** | **0.013** | **0.057** | **2213.434** | **0.227** | **1.000** |
| H,-1 - H+,0 | 0.050 | 0.088 | 107.926 | 0.563 | 1.000 |
| **H,-1 - H,1** | **-0.142** | **0.051** | **2217.319** | **-2.765** | **0.149** |
| H,-1 - H+,1 | -0.037 | 0.087 | 103.187 | -0.423 | 1.000 |
| **H,-1 - H,2** | **-0.277** | **0.058** | **2229.053** | **-4.791** | **<0.001*** ** |
| H,-1 - H+,2 | -0.069 | 0.090 | 116.076 | -0.765 | 0.999 |
| **H,-1 - H,3** | **-0.222** | **0.058** | **2234.701** | **-3.852** | **0.005**** |
| H,-1 - H+,3 | 0.006 | 0.089 | 109.947 | 0.066 | 1.000 |
| H+,-1 - H,0 | -0.178 | 0.089 | 110.503 | -2.005 | 0.597 |
| **H+,-1 - H+,0** | **-0.141** | **0.048** | **2211.834** | **-2.947** | **0.094** |
| H+,-1 - H,1 | -0.333 | 0.084 | 90.422 | -3.942 | 0.006** |
| **H+,-1 - H+,1** | **-0.228** | **0.046** | **2208.007** | **-4.976** | **<0.001*** ** |
| H+,-1 - H,2 | -0.469 | 0.088 | 105.757 | -5.342 | <0.001*** |

| **H,-1 - H,2** | **-0.273** | **0.055** | **2226.726** | **-4.983** | **<0.001*****
| H,-1 - C,3 | -0.204 | 0.053 | 2227.487 | -3.835 | 0.005** |
| **H,-1 - H,3** | **-0.195** | **0.054** | **2236.850** | **-3.592** | **0.012*** |
| **C,0 - H,0** | **-0.390** | **0.047** | **2211.157** | **-8.290** | **<0.001***** |
| **C,0 - C,1** | **-0.310** | **0.042** | **2207.577** | **-7.321** | **<0.001***** |
| C,0 - H,1 | -0.321 | 0.041 | 2230.690 | -7.774 | <0.001*** |
| **C,0 - C,2** | **-0.391** | **0.048** | **2219.303** | **-8.097** | **<0.001***** |
| C,0 - H,2 | -0.408 | 0.049 | 2232.506 | -8.304 | <0.001*** |
| **C,0 - C,3** | **-0.339** | **0.046** | **2220.125** | **-7.325** | **<0.001***** |
| C,0 - H,3 | -0.330 | 0.048 | 2242.830 | -6.858 | <0.001*** |
| H,0 - C,1 | 0.080 | 0.044 | 2221.179 | 1.833 | 0.714 |
| **H,0 - H,1** | **0.069** | **0.041** | **2227.913** | **1.660** | **0.818** |
| H,0 - C,2 | -0.001 | 0.049 | 2223.051 | -0.019 | 1.000 |
| **H,0 - H,2** | **-0.018** | **0.049** | **2227.028** | **-0.368** | **1.000** |
| H,0 - C,3 | 0.051 | 0.047 | 2225.491 | 1.093 | 0.985 |
| **H,0 - H,3** | **0.060** | **0.048** | **2240.114** | **1.243** | **0.965** |
| **C,1 - H,1** | **-0.011** | **0.037** | **2237.103** | **-0.302** | **1.000** |
| **C,1 - C,2** | **-0.081** | **0.044** | **2212.857** | **-1.831** | **0.716** |
| C,1 - H,2 | -0.098 | 0.046 | 2238.764 | -2.141 | 0.498 |
| **C,1 - C,3** | **-0.029** | **0.042** | **2221.238** | **-0.681** | **1.000** |
| C,1 - H,3 | -0.020 | 0.044 | 2245.906 | -0.446 | 1.000 |
| H,1 - C,2 | -0.070 | 0.043 | 2240.324 | -1.608 | 0.844 |
| **H,1 - H,2** | **-0.087** | **0.042** | **2215.873** | **-2.076** | **0.545** |
| H,1 - C,3 | -0.018 | 0.041 | 2244.356 | -0.426 | 1.000 |
| **H,1 - H,3** | **-0.009** | **0.041** | **2233.590** | **-0.211** | **1.000** |
| **C,2 - H,2** | **-0.017** | **0.051** | **2239.442** | **-0.336** | **1.000** |
| **C,2 - C,3** | **0.052** | **0.047** | **2209.017** | **1.110** | **0.984** |
| C,2 - H,3 | 0.061 | 0.050 | 2246.484 | 1.221 | 0.969 |
| H,2 - C,3 | 0.069 | 0.049 | 2242.411 | 1.405 | 0.926 |
| **H,2 - H,3** | **0.078** | **0.048** | **2219.511** | **1.634** | **0.831** |
| **C,3 - H,3** | **0.009** | **0.048** | **2246.911** | **0.187** | **1.000** |
| Partner:Treatment interaction | | - | - | - | - |
| **C,H - H,H** | **-0.136** | **0.036** | **2238.191** | **-3.803** | **0.001**** |
| **C,H - C,H+** | **0.101** | **0.077** | **63.411** | **1.311** | **0.559** |
| C,H - H,H+ | 0.070 | 0.077 | 62.477 | 0.912 | 0.798 |

| | | | | | |
|---|---|---|---|---|---|
| **H+,-1 - H+,2** | **-0.260** | **0.051** | **2210.669** | **-5.087** | **<0.001***** |
| H+,-1 - H,3 | -0.413 | 0.087 | 102.286 | -4.763 | <0.001*** |
| **H+,-1 - H+,3** | **-0.185** | **0.049** | **2213.912** | **-3.751** | **0.007**** |
| **H,0 - H+,0** | **0.037** | **0.085** | **92.223** | **0.433** | **1.000** |
| **H,0 - H,1** | **-0.155** | **0.046** | **2223.429** | **-3.383** | **0.025*** |
| H,0 - H+,1 | -0.050 | 0.084 | 87.833 | -0.595 | 1.000 |
| **H,0 - H,2** | **-0.290** | **0.053** | **2227.936** | **-5.520** | **<0.001***** |
| H,0 - H+,2 | -0.082 | 0.086 | 99.763 | -0.945 | 0.994 |
| **H,0 - H,3** | **-0.235** | **0.052** | **2235.931** | **-4.501** | **<0.001***** |
| H,0 - H+,3 | -0.007 | 0.085 | 94.035 | -0.084 | 1.000 |
| H+,0 - H,1 | -0.191 | 0.080 | 73.783 | -2.387 | 0.349 |
| **H+,0 - H+,1** | **-0.087** | **0.038** | **2219.435** | **-2.270** | **0.409** |
| H+,0 - H,2 | -0.327 | 0.084 | 87.806 | -3.912 | 0.007** |
| **H+,0 - H+,2** | **-0.118** | **0.044** | **2215.176** | **-2.689** | **0.179** |
| H+,0 - H,3 | -0.271 | 0.083 | 84.400 | -3.289 | 0.045* |
| **H+,0 - H+,3** | **-0.044** | **0.042** | **2210.750** | **-1.052** | **0.989** |
| **H,1 - H+,1** | **0.105** | **0.079** | **69.890** | **1.323** | **0.945** |
| **H,1 - H,2** | **-0.136** | **0.045** | **2221.435** | **-3.051** | **0.070** |
| H,1 - H+,2 | 0.073 | 0.082 | 80.634 | 0.890 | 0.996 |
| **H,1 - H,3** | **-0.080** | **0.044** | **2231.189** | **-1.830** | **0.716** |
| H,1 - H+,3 | 0.147 | 0.080 | 75.439 | 1.832 | 0.713 |
| H+,1 - H,2 | -0.241 | 0.083 | 83.508 | -2.911 | 0.119 |
| **H+,1 - H+,2** | **-0.032** | **0.042** | **2210.220** | **-0.766** | **0.999** |
| H+,1 - H,3 | -0.185 | 0.082 | 80.281 | -2.266 | 0.422 |
| **H+,1 - H+,3** | **0.043** | **0.039** | **2214.573** | **1.087** | **0.986** |
| **H,2 - H+,2** | **0.209** | **0.085** | **95.185** | **2.448** | **0.310** |
| **H,2 - H,3** | **0.056** | **0.050** | **2215.502** | **1.118** | **0.983** |
| H,2 - H+,3 | 0.283 | 0.084 | 89.562 | 3.375 | 0.035* |
| H+,2 - H,3 | -0.153 | 0.084 | 91.950 | -1.816 | 0.723 |
| **H+,2 - H+,3** | **0.075** | **0.045** | **2210.373** | **1.663** | **0.816** |
| **H,3 - H+,3** | **0.228** | **0.083** | **86.395** | **2.745** | **0.173** |
| Partner:Batch interaction | | | | | |
| **C,1 - H,1** | **-0.132** | **0.031** | **2241.341** | **-4.329** | **<0.001***** |
| **C,1 - C,2** | **-0.310** | **0.077** | **61.934** | **-4.033** | **0.001**** |
| C,1 - H,2 | -0.344 | 0.077 | 62.924 | -4.488 | <0.001*** |

| | | | | | | | | | | | |
|---|---|---|---|---|---|---|---|---|---|---|---|
| H,H - C,H+ | 0.237 | 0.077 | 62.634 | 3.097 | 0.015* | H,1 - C,2 | -0.177 | 0.076 | 60.532 | -2.324 | 0.104 |
| **H,H - H,H+** | **0.206** | **0.077** | **62.478** | **2.691** | **0.044*** | **H,1 - H,2** | **-0.212** | **0.076** | **60.360** | **-2.795** | **0.034*** |
| **C,H+ - H,H+** | **-0.031** | **0.030** | **2246.941** | **-1.021** | **0.737** | **C,2 - H,2** | **-0.035** | **0.034** | **2246.804** | **-1.020** | **0.738** |
| <u>Treatment:Batch interaction</u> | | | | | | | | | | | |
| **H,1 - H+,1** | **0.011** | **0.100** | **48.467** | **0.108** | **1.000** | | | | | | |
| **H,1 - H,2** | **-0.404** | **0.105** | **54.967** | **-3.831** | **0.002**** | | | | | | |
| H,1 - H+,2 | -0.107 | 0.098 | 45.975 | -1.095 | 0.694 | | | | | | |
| H+,1 - H,2 | -0.415 | 0.109 | 56.833 | -3.822 | 0.002** | | | | | | |
| **H+,1 - H+,2** | **-0.118** | **0.101** | **47.419** | **-1.171** | **0.648** | | | | | | |
| **H,2 - H+,2** | **0.297** | **0.107** | **54.466** | **2.780** | **0.036*** | | | | | | |

DATA COMPOSITION TABLES

**Table S3: Acoustic values for acoustic proxies per group (Treatment: Partner: Phase of trial)**. The number of vocalisations per group is indicated and mean, standard deviation (sd), standard error (se) and 95% confidence interval (ci) are indicated for the vocalisation duration and all the spectral parameters used to build the spectral acoustic score (LD1)

| | | | | Grunt duration (s) | | | | Mean (Hz) | | | | Median (Hz) | | | | Mode (Hz) | | | |
|---|---|---|---|---|---|---|---|---|---|---|---|---|---|---|---|---|---|---|---|
| Treatment | Partner | Phase of trial | N | mean | sd | se | ci | mean | sd | se | ci | mean | sd | se | ci | mean | sd | se | ci |
| H | C | -1 | 56 | 0.270 | 0.150 | 0.020 | 0.040 | 1684.124 | 327.780 | 43.801 | 87.780 | 620.722 | 348.976 | 46.634 | 93.456 | 311.146 | 43.253 | 5.780 | 11.583 |
| H | C | 0 | 86 | 0.230 | 0.166 | 0.018 | 0.036 | 1567.744 | 269.436 | 29.054 | 57.767 | 537.490 | 194.757 | 21.001 | 41.756 | 321.186 | 43.040 | 4.641 | 9.228 |
| H | C | 1 | 158 | 0.378 | 0.238 | 0.019 | 0.037 | 1603.305 | 292.058 | 23.235 | 45.893 | 515.844 | 182.099 | 14.487 | 28.615 | 301.127 | 42.061 | 3.346 | 6.609 |
| H | C | 2 | 94 | 0.437 | 0.242 | 0.025 | 0.050 | 1507.209 | 245.191 | 25.289 | 50.220 | 462.287 | 134.290 | 13.851 | 27.505 | 293.664 | 42.098 | 4.342 | 8.623 |
| H | C | 3 | 107 | 0.427 | 0.250 | 0.024 | 0.048 | 1525.283 | 294.430 | 28.464 | 56.432 | 517.038 | 231.680 | 22.397 | 44.405 | 298.408 | 41.811 | 4.042 | 8.014 |
| H | H | -1 | 63 | 0.306 | 0.185 | 0.023 | 0.047 | 1739.732 | 311.840 | 39.288 | 78.536 | 590.546 | 241.963 | 30.485 | 60.938 | 328.205 | 41.924 | 5.282 | 10.558 |
| H | H | 0 | 78 | 0.369 | 0.193 | 0.022 | 0.044 | 1653.975 | 305.643 | 34.607 | 68.912 | 554.748 | 249.115 | 28.207 | 56.167 | 300.592 | 50.770 | 5.749 | 11.447 |
| H | H | 1 | 183 | 0.389 | 0.195 | 0.014 | 0.028 | 1602.807 | 286.368 | 21.169 | 41.768 | 534.871 | 227.058 | 16.785 | 33.117 | 295.346 | 42.871 | 3.169 | 6.253 |
| H | H | 2 | 81 | 0.404 | 0.184 | 0.020 | 0.041 | 1618.200 | 292.162 | 32.462 | 64.602 | 563.397 | 240.758 | 26.751 | 53.236 | 297.924 | 43.314 | 4.813 | 9.578 |
| H | H | 3 | 83 | 0.408 | 0.181 | 0.020 | 0.039 | 1519.422 | 270.347 | 29.674 | 59.032 | 503.429 | 182.851 | 20.071 | 39.927 | 294.347 | 42.547 | 4.670 | 9.291 |
| H+ | C | -1 | 82 | 0.259 | 0.120 | 0.013 | 0.026 | 1502.188 | 264.115 | 29.167 | 58.032 | 472.045 | 129.253 | 14.274 | 28.400 | 310.480 | 49.324 | 5.447 | 10.838 |
| H+ | C | 0 | 139 | 0.246 | 0.116 | 0.010 | 0.019 | 1555.290 | 231.077 | 19.600 | 38.755 | 496.166 | 146.289 | 12.408 | 24.534 | 315.408 | 43.770 | 3.713 | 7.341 |
| H+ | C | 1 | 146 | 0.324 | 0.142 | 0.012 | 0.023 | 1587.505 | 284.500 | 23.545 | 46.537 | 503.989 | 174.373 | 14.431 | 28.523 | 302.514 | 45.484 | 3.764 | 7.440 |
| H+ | C | 2 | 93 | 0.345 | 0.174 | 0.018 | 0.036 | 1497.941 | 241.367 | 25.029 | 49.709 | 479.074 | 163.642 | 16.969 | 33.702 | 296.065 | 43.556 | 4.517 | 8.970 |
| H+ | C | 3 | 113 | 0.298 | 0.151 | 0.014 | 0.028 | 1507.152 | 292.735 | 27.538 | 54.563 | 501.177 | 191.904 | 18.053 | 35.769 | 304.580 | 44.377 | 4.175 | 8.271 |
| H+ | H | -1 | 70 | 0.243 | 0.112 | 0.013 | 0.027 | 1575.042 | 244.688 | 29.246 | 58.344 | 498.713 | 203.349 | 24.305 | 48.487 | 303.925 | 45.780 | 5.472 | 10.916 |
| H+ | H | 0 | 142 | 0.327 | 0.140 | 0.012 | 0.023 | 1555.213 | 265.097 | 22.246 | 43.980 | 496.022 | 168.004 | 14.099 | 27.872 | 299.396 | 48.115 | 4.038 | 7.982 |
| H+ | H | 1 | 247 | 0.334 | 0.157 | 0.010 | 0.020 | 1596.026 | 282.566 | 17.979 | 35.413 | 506.687 | 175.430 | 11.162 | 21.986 | 303.742 | 47.220 | 3.005 | 5.918 |
| H+ | H | 2 | 113 | 0.357 | 0.183 | 0.017 | 0.034 | 1594.982 | 307.845 | 28.960 | 57.380 | 555.247 | 235.247 | 22.130 | 43.848 | 293.222 | 47.574 | 4.475 | 8.867 |
| H+ | H | 3 | 136 | 0.336 | 0.182 | 0.016 | 0.031 | 1581.890 | 276.006 | 23.667 | 46.807 | 514.092 | 190.153 | 16.305 | 32.247 | 301.681 | 43.880 | 3.763 | 7.442 |

| | | | | First Quartile Q25 (Hz) | | | | Third Quartile Q75 (Hz) | | | | Centroid (Hz) | | | |
|---|---|---|---|---|---|---|---|---|---|---|---|---|---|---|---|
| Treatment | Partner | Phase of test | N | mean | sd | se | ci | mean | sd | se | ci | mean | sd | se | ci |
| H | C | -1 | 56 | 302.041 | 54.385 | 7.267 | 14.564 | 2420.879 | 819.325 | 109.487 | 219.417 | 1684.124 | 327.780 | 43.801 | 87.780 |

| | | | | | | | | | | | | | | | |
|---|---|---|---|---|---|---|---|---|---|---|---|---|---|---|---|
| H | C | 0 | 86 | 306.670 | 45.937 | 4.954 | 9.849 | 2027.396 | 699.634 | 75.443 | 150.002 | 1567.744 | 269.436 | 29.054 | 57.767 |
| H | C | 1 | 158 | 286.549 | 40.281 | 3.205 | 6.330 | 2231.075 | 753.146 | 59.917 | 118.348 | 1603.305 | 292.058 | 23.235 | 45.893 |
| H | C | 2 | 94 | 274.024 | 30.444 | 3.140 | 6.235 | 1965.396 | 663.033 | 68.387 | 135.802 | 1507.209 | 245.191 | 25.289 | 50.220 |
| H | C | 3 | 107 | 282.058 | 37.747 | 3.649 | 7.235 | 1975.064 | 778.304 | 75.241 | 149.174 | 1525.283 | 294.430 | 28.464 | 56.432 |
| H | H | -1 | 63 | 310.344 | 47.490 | 5.983 | 11.960 | 2565.358 | 763.230 | 96.158 | 192.217 | 1739.732 | 311.840 | 39.288 | 78.536 |
| H | H | 0 | 78 | 284.813 | 38.991 | 4.415 | 8.791 | 2302.733 | 750.096 | 84.932 | 169.121 | 1653.975 | 305.643 | 34.607 | 68.912 |
| H | H | 1 | 183 | 287.708 | 41.677 | 3.081 | 6.079 | 2197.610 | 731.375 | 54.065 | 106.674 | 1602.807 | 286.368 | 21.169 | 41.768 |
| H | H | 2 | 81 | 287.179 | 40.046 | 4.450 | 8.855 | 2219.726 | 695.209 | 77.245 | 153.723 | 1618.200 | 292.162 | 32.462 | 64.602 |
| H | H | 3 | 83 | 280.154 | 33.959 | 3.727 | 7.415 | 1974.527 | 661.681 | 72.629 | 144.482 | 1519.422 | 270.347 | 29.674 | 59.032 |
| H+ | C | -1 | 82 | 284.428 | 37.511 | 4.142 | 8.242 | 1986.813 | 765.804 | 84.569 | 168.266 | 1502.188 | 264.115 | 29.167 | 58.032 |
| H+ | C | 0 | 139 | 288.908 | 42.600 | 3.613 | 7.145 | 2114.444 | 645.644 | 54.763 | 108.283 | 1555.290 | 231.077 | 19.600 | 38.755 |
| H+ | C | 1 | 146 | 286.076 | 40.886 | 3.384 | 6.688 | 2213.157 | 736.747 | 60.974 | 120.512 | 1587.505 | 284.500 | 23.545 | 46.537 |
| H+ | C | 2 | 93 | 272.717 | 30.536 | 3.166 | 6.289 | 1991.540 | 649.380 | 67.338 | 133.738 | 1497.941 | 241.367 | 25.029 | 49.709 |
| H+ | C | 3 | 113 | 286.920 | 45.241 | 4.256 | 8.433 | 1966.975 | 783.150 | 73.673 | 145.973 | 1507.152 | 292.735 | 27.538 | 54.563 |
| H+ | H | -1 | 70 | 279.435 | 37.265 | 4.454 | 8.885 | 2240.502 | 620.204 | 74.129 | 147.882 | 1575.042 | 244.688 | 29.246 | 58.344 |
| H+ | H | 0 | 142 | 282.495 | 39.513 | 3.316 | 6.555 | 2087.726 | 738.101 | 61.940 | 122.451 | 1555.213 | 265.097 | 22.246 | 43.980 |
| H+ | H | 1 | 247 | 285.056 | 39.453 | 2.510 | 4.944 | 2240.406 | 729.732 | 46.432 | 91.454 | 1596.026 | 282.566 | 17.979 | 35.413 |
| H+ | H | 2 | 113 | 287.465 | 44.640 | 4.199 | 8.320 | 2214.684 | 774.470 | 72.856 | 144.355 | 1594.982 | 307.845 | 28.960 | 57.380 |
| H+ | H | 3 | 136 | 280.155 | 34.604 | 2.967 | 5.868 | 2203.555 | 737.813 | 63.267 | 125.122 | 1581.890 | 276.006 | 23.667 | 46.807 |

| | | | | **Sh** | | | | **Sfm** | | | | **Enthropy** | | | |
|---|---|---|---|---|---|---|---|---|---|---|---|---|---|---|---|
| **Treatment** | **Partner** | **Phase of trial** | **N** | **mean** | **sd** | **se** | **ci** | **mean** | **sd** | **se** | **ci** | **mean** | **sd** | **se** | **ci** |
| H | C | -1 | 56 | 0.790 | 0.059 | 0.008 | 0.016 | 0.503 | 0.101 | 0.014 | 0.027 | 0.600 | 0.045 | 0.006 | 0.012 |
| H | C | 0 | 86 | 0.772 | 0.051 | 0.006 | 0.011 | 0.462 | 0.083 | 0.009 | 0.018 | 0.579 | 0.039 | 0.004 | 0.008 |
| H | C | 1 | 158 | 0.777 | 0.051 | 0.004 | 0.008 | 0.479 | 0.089 | 0.007 | 0.014 | 0.593 | 0.039 | 0.003 | 0.006 |
| H | C | 2 | 94 | 0.760 | 0.050 | 0.005 | 0.010 | 0.452 | 0.080 | 0.008 | 0.016 | 0.585 | 0.035 | 0.004 | 0.007 |
| H | C | 3 | 107 | 0.764 | 0.059 | 0.006 | 0.011 | 0.456 | 0.091 | 0.009 | 0.017 | 0.587 | 0.045 | 0.004 | 0.009 |
| H | H | -1 | 63 | 0.794 | 0.052 | 0.007 | 0.013 | 0.509 | 0.088 | 0.011 | 0.022 | 0.603 | 0.042 | 0.005 | 0.011 |
| H | H | 0 | 78 | 0.782 | 0.056 | 0.006 | 0.013 | 0.491 | 0.092 | 0.010 | 0.021 | 0.600 | 0.043 | 0.005 | 0.010 |
| H | H | 1 | 183 | 0.778 | 0.053 | 0.004 | 0.008 | 0.480 | 0.088 | 0.007 | 0.013 | 0.596 | 0.041 | 0.003 | 0.006 |
| H | H | 2 | 81 | 0.785 | 0.054 | 0.006 | 0.012 | 0.488 | 0.090 | 0.010 | 0.020 | 0.604 | 0.040 | 0.004 | 0.009 |
| H | H | 3 | 83 | 0.766 | 0.056 | 0.006 | 0.012 | 0.458 | 0.087 | 0.010 | 0.019 | 0.591 | 0.044 | 0.005 | 0.010 |
| H+ | C | -1 | 82 | 0.759 | 0.049 | 0.005 | 0.011 | 0.447 | 0.080 | 0.009 | 0.018 | 0.575 | 0.036 | 0.004 | 0.008 |
| H+ | C | 0 | 139 | 0.770 | 0.045 | 0.004 | 0.008 | 0.464 | 0.075 | 0.006 | 0.013 | 0.580 | 0.039 | 0.003 | 0.007 |

| | | | | | | | | | | | | | | | |
|---|---|---|---|---|---|---|---|---|---|---|---|---|---|---|---|
| H+ | C | 1 | 146 | 0.775 | 0.049 | 0.004 | 0.008 | 0.475 | 0.084 | 0.007 | 0.014 | 0.592 | 0.038 | 0.003 | 0.006 |
| H+ | C | 2 | 93 | 0.762 | 0.049 | 0.005 | 0.010 | 0.452 | 0.075 | 0.008 | 0.016 | 0.583 | 0.039 | 0.004 | 0.008 |
| H+ | C | 3 | 113 | 0.761 | 0.057 | 0.005 | 0.011 | 0.451 | 0.088 | 0.008 | 0.016 | 0.579 | 0.043 | 0.004 | 0.008 |
| H+ | H | -1 | 70 | 0.772 | 0.045 | 0.005 | 0.011 | 0.472 | 0.076 | 0.009 | 0.018 | 0.584 | 0.033 | 0.004 | 0.008 |
| H+ | H | 0 | 142 | 0.768 | 0.054 | 0.005 | 0.009 | 0.466 | 0.082 | 0.007 | 0.014 | 0.588 | 0.039 | 0.003 | 0.007 |
| H+ | H | 1 | 247 | 0.776 | 0.051 | 0.003 | 0.006 | 0.479 | 0.085 | 0.005 | 0.011 | 0.593 | 0.040 | 0.003 | 0.005 |
| H+ | H | 2 | 113 | 0.782 | 0.056 | 0.005 | 0.010 | 0.481 | 0.094 | 0.009 | 0.017 | 0.598 | 0.044 | 0.004 | 0.008 |
| H+ | H | 3 | 136 | 0.776 | 0.050 | 0.004 | 0.009 | 0.475 | 0.083 | 0.007 | 0.014 | 0.591 | 0.041 | 0.004 | 0.007 |
